# Mimicking tumor cell heterogeneity of colorectal cancer in a patient-derived organoid-fibroblast model

**DOI:** 10.1101/2022.03.07.483214

**Authors:** Velina S Atanasova, Crhistian de Jesus Cardona, Vaclav Hejret, Andreas Tiefenbacher, Loan Tran, Carina Binder, Theresia Mair, Julijan Kabiljo, Janik Clement, Katharina Woeran, Barbara Neudert, Markus Hengstschläger, Markus Mitterhauser, Leonhard Müllauer, Boris Tichy, Michael Bergmann, Gabriele Schweikert, Markus Hartl, Helmut Dolznig, Gerda Egger

## Abstract

Patient-derived organoid (PDO) cancer models are generated from epithelial tumor cells. Although they reflect the molecular tumor characteristics, they lack the complexity of the tumor microenvironment, which is a key driver of tumorigenesis and therapy response. Here, we present a colorectal cancer (CRC) organoid model that incorporates epithelial cells and stromal fibroblasts from the same patient. Molecular characterization of primary cancer associated fibroblasts (CAFs) and matched normal fibroblasts (NF) revealed proteomic, secretome and gene expression differences in pathways associated with tumor related fibroblast function. Further, CAFs retained higher motility compared to NFs *in vitro*. Importantly, both CAFs and NFs supported cancer cell proliferation in 3D co-cultures, without the addition of classical niche factors. PDOs grown together with fibroblasts displayed a larger cellular heterogeneity of tumor cells compared to mono-cultures, and closely resembled the *in vivo* tumor morphology. This was also confirmed by the calculation of cellular proportions of epithelial cell subtypes in organoid mono-versus co-cultures, which were inferred through bioinformatics deconvolution of bulk RNA sequencing data using published single cell RNA sequencing datasets from CRC tissues. Additionally, we observed a mutual crosstalk between tumor cells and fibroblasts in the co-cultures. This was manifested by majorly deregulated pathways such as cell-cell communication and extracellular matrix remodeling in the organoids. For the fibroblasts, we observed enhanced expression of tumor induced marker genes and cytokines characteristic for myo- and immunogenic fibroblasts. This model will be vital as a physiological personalized tumor model to study disease mechanisms and therapy response in CRC.

**One Sentence Summary:** Patient matched fibroblasts support tumor organoid growth in 3D co-culture and maintain intratumoral cellular heterogeneity and histo-morphology.

## INTRODUCTION

Colorectal cancer (CRC) remains a main health burden worldwide, being the 4^th^ leading cause of cancer-related death (*1*). Although there has been great scientific advance, patient treatment options for advanced CRC remain limited and face poor response prediction. The standard of care therapy remains unchanged since decades and predominantly relies on chemotherapy, often associated with chemoresistance (*2*).

The recent development of advanced tissue culture models, derived from normal as well as tumor tissues, has given scientists an indispensable tool to work with models reflecting the *in vivo* tumor characteristics. These so-called patient-derived organoids (PDOs) are promising tools for drug screening and have been shown to predict patient response for several different tumor entities (*3, 4*). However, CRC organoids generated from metastatic tumors could not predict therapy response of patients for combination therapies with 5-fluorouracil (5-FU) plus oxaliplatin (*5, 6*). This might be due to the specific *in vitro* culture conditions or the lack of the complex tumor microenvironment (TME) in PDOs. The TME represents the non-malignant part of the tumor and includes the infiltrating immune cells, endothelial cells, nerves and fibroblasts, often referred to as cancer associated fibroblasts (CAFs). In fact, the tumor surrounding cells can impact not only on the tumor development but have a large role in therapy response and resistance. Patients with tumors displaying high CAF signatures face lower rates of survival with high statistical prediction (*7, 8*). The consensus molecular subtypes (CMS) of CRC include a mesenchymal subtype CMS4, which is characterized by stromal infiltration, activation of TGFβ signaling and genes involved in epithelial-to-mesenchymal transition (EMT), angiogenesis, matrix remodeling and complement-mediated inflammatory system. Furthermore, CMS4 tumors often display with metastases and thus poorer patient survival compared to the other subtypes (*9*).

Fibroblasts are the main cell type in the stroma and its main remodelers, as well as a major source of cytokines, growth and survival factors. While fibroblasts in healthy tissues become activated only upon tissue damage such as wounding and trigger a program of tissue repair until the wound has healed, CAFs are characterized by a constantly activated ‘wound healing’ phenotype and tumor supportive properties (*10*). Compared to normal fibroblasts (NFs), CAFs exhibit increased proliferation, secretion of extracellular matrix and a specific cytokine profile (*11*). CAFs represent a heterogeneous and plastic population of cells that can be derived from diverse cellular resources including resident fibroblasts, endothelial cells, adipocytes and others. Recent single cell RNA sequencing (scRNA-seq) efforts of CRC tumors revealed two main CAF populations including inflammatory (iCAFs) and myofibroblast-like (myCAFs) fibroblasts, which are involved in immune-crosstalk and matrix remodeling, respectively (*12–14*).

Even though specific CAF markers remain elusive, the clinical relevance of CAFs in tumors is indisputable – the abundance of CAFs within a tumor has been associated with poor prognosis in cancer types such as colon (*15, 16*), breast (*17*) and pancreas (*18*). Furthermore, there is an extensive cross talk between fibroblasts and other stromal cell types including immune cells, where CAFs exhibit immune suppressive properties thus preventing tumor recognition by the immune system (*19, 20*). In fact, early co-culture experiments have demonstrated that irradiated fibroblasts better support tumor cell growth compared to nonirradiated fibroblasts pointing out that these cells actively affect the tumor and can potentially affect response to therapy (*10, 21–24*). Further, a recent report identified an association of iCAFs with poor prognosis in rectal cancer following irradiation (*25*).

Here, we describe the establishment and characterization of an advanced co-culture PDO model, where CRC derived epithelial cells and patient-matched fibroblasts can be cocultured. With this model, we studied the cross-talk between tumor organoids and normal or cancer fibroblasts from the same patient and the expression alterations the two cell types induce upon one another. Importantly, this model allows for the cultivation of PDOs without the addition of niche factors such as EGF, R-spondin1, Noggin, Wnt3a and inhibitors such as SB202190, A8301, which are known to affect fibroblast phenotype and proliferation (*26, 27*). We find that PDOs co-cultured with fibroblasts show a heterogenous cellular composition and closer resemblance to primary tumor tissues based on histomorphology and gene expression profiles inferred from scRNA-seq data using a bioinformatics deconvolution strategy. Overall, this system represents a valuable tool for drug development, precision medicine, therapy-response prediction and improved understanding of the carcinoma biology in respect to tumor-stroma crosstalk.

## RESULTS

### Isolation of primary cells and generation of the co-culture model

In order to generate a physiological tumoroid model, which also contains stromal fibroblasts, we isolated epithelial tumor cells and fibroblasts from the same patient (Fig. 1A). Fibroblasts were cultivated in 2-D and derived both from the tumor as well as normal adjacent tissues. Organoids were generated in Matrigel following standard protocols (*28*). For co-cultivation of the different cell types, we developed three individual setups using matched tumor and fibroblast cells from three individual patients with moderately differentiated tumors (G2), carrying *APC* mutations, and located in the right sided colon (P1-3, table S1). Both tumor cells and fibroblasts were either cultivated together in Matrigel plugs, or tumor cells and fibroblasts were divided into individual layers in air/liquid interphase cultures. For these separated cultures, fibroblasts were grown in collagen and organoids seeded on top directly or in Matrigel separated by a 0.45 μm membrane on a transwell insert.

**Fig. 1:**
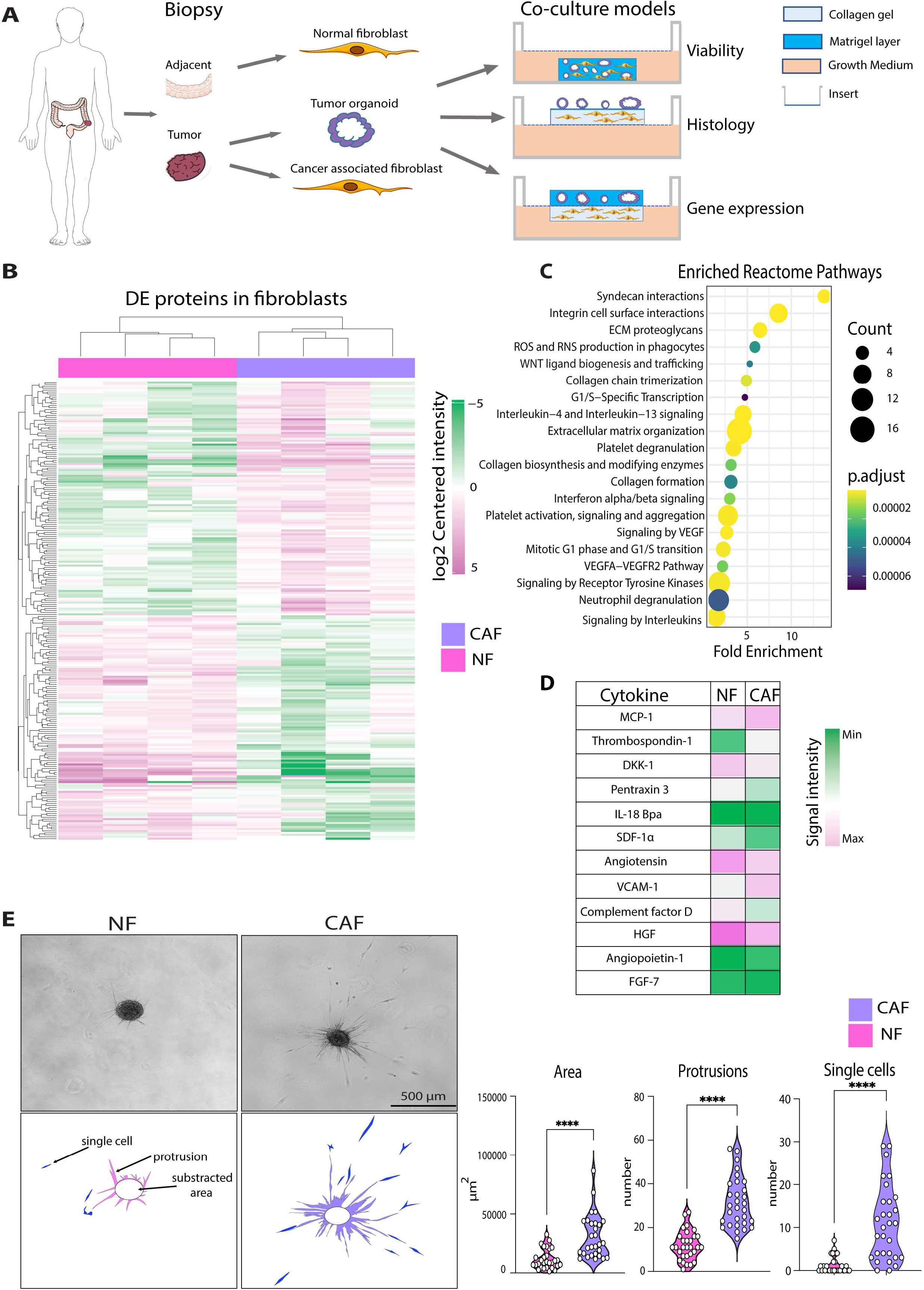
Molecular and phenotypic differences of NFs and CAFs. **(A)** Normal (NFs) and cancer associated fibroblasts (CAFs) as well as patient derived organoids (PDOs) are isolated from colon tumors or normal adjacent mucosa. PDOs together with either NFs or CAFs are cultured in different co-culture models containing either Matrigel alone or with collagen I as supportive matrices. Individual setups were used for molecular analyses as indicated (right). **(B)** Hierarchical clustering of significantly differentially expressed proteins between 4 individual pairs of NFs and CAFs as analyzed by label free MS/MS (*P* < 0.05, absolute log2 fold change ≥1) **(C)** Bubble chart describes 20 of the top significant pathways detected by Reactome pathway analysis of differentially expressed proteins. The size of the circles corresponds to the number of proteins changed in the respective pathway. Adjusted *P* values are indicated by different colors according to the color gradient **(D)** Intensity values of significantly differentially secreted proteins between 7 matched NF and CAF pairs as analyzed by human proteome profiler assays (*P* < 0.05). **(E)** Representative microscopic pictures of NF and CAF spheroids embedded into collagen I for 24h (left). Illustration of quantification of the sprouting area, number of protrusions and single cells in CAFs versus NFs (lower image) Violin plots show measurements of the sprouting area (total area minus spheroid core area), number of protrusions and number of single cells detached from the spheroids (right). Experiments were performed in triplicates and 10 individual spheroids were analyzed for each experiment (*N* = 30). \**P* < 0.05, \*\**P* < 0.005, \*\*\**P* < 0.001. Two-tailed unpaired Student’s *t* test.

### Characterization of normal and tumor stromal fibroblasts by proteomic and secretome analysis

As stromal fibroblasts in normal tissues *in vivo* exhibit multiple differences compared to CAFs, we first investigated whether these variances remain stable between primary cultured NFs and CAFs isolated from adjacent normal tissue and CRC specimen. For this, we collected total protein from patient-matched NFs and CAFs in early passages (3-5). By label free MS/MS proteomic analysis of four independent patient-matched NF and CAF pairs, a total of 4796 protein groups were detected out of which 235 were significantly differentially expressed (*P*<0.05, absolute log2 fold change ≥1) (Fig.1B and table S2). Reactome pathway analysis revealed that molecules related to extracellular matrix organization, collagen biosynthesis and cell attachment were deregulated in CAFs compared to NFs (Fig. 1C and fig. S1A). Additionally, several immune system pathways such as interleukin or interferon signaling were significantly affected by differentially expressed proteins between the two fibroblast groups. Among the top differentially expressed proteins between NFs and CAFs we found several proteins related to cell migration and cell contraction including hyaluronan-binding protein, nestin, myosin light chain kinase, Rho-related GTP-binding protein RhoE and supervillin, which were increased 2 to 3-fold in CAFs compared to NFs (table S2).

Secretome analysis of conditioned media of seven pairs of matched NFs and CAFs using cytokine arrays identified twelve molecules that were significantly deregulated between CAFs and NFs (Fig. 1D, fig. S1B). Among the top differentially secreted molecules were MCP1 (CCL2) and Thrombospondin 1 (both increased in CAFs), as well as DKK1 and Pentraxin 3 (both increased in NFs). MCP-1 is a promoter of lung metastasis in breast cancer by promoting angiogenesis (*29*) while Thrombospondin 1 is a TGFβ activator and has been found overexpressed in tumor stroma (*30*). DKK1 is an antagonist of Wnt signaling and being downregulated at the offset of adeno-carcinoma formation (*31*). Finally, the exact role of Pentraxin 3 in cancer has not been fully elucidated, however, it was suggested as an oncosuppressor in mice and human via regulation of Complement-dependent tumor-promoting inflammation (*32*). Together, these data show that NFs and CAFs retain their functional proteome and secretome *in vitro*, which reflects tumor associated pathways in CAFs.

### Collagen motility assay reveals phenotypic differences between CAFs and NFs

Based on the proteome and secretome data that pointed towards deregulated pathways in extracellular matrix remodeling and thus a possible difference in cell motility we evaluated whether these molecular changes translate into phenotypic differences between NFs and CAFs. Indeed, 3D collagen gel motility assays of fibroblast spheroids (*33*) revealed significantly higher spread areas, numbers of protrusions and numbers of single cells detached from the spheroid structures (Fig. 1E and fig. S1C,D) in three matched early passage (2-5) NF and CAF pairs. These findings suggest that the matrix protein alterations found in the proteomic analyses indeed led to a phenotypic difference manifested by increased motility of CAFs in collagen gels.

### Both normal and cancer associated fibroblasts support organoid growth in a co-culture model

Following NF and CAF characterization, we sought to investigate the crosstalk between tumor cells and stromal fibroblasts. For this purpose, we established a model where fibroblasts and PDOs derived from the same patient were co-cultured in Matrigel drops in minimal medium (1% FCS). We omitted the addition of any niche factors or inhibitors that are usually added to conventional organoid growth medium to allow for expansion of stemlike cells and inhibition of differentiation signals. Importantly, these factors were shown to alter fibroblast signaling and phenotype (*34, 35*). Particularly, inhibitors of TGFβ signaling (A8301) or cell cycle (SB202190) (*36*) would potentially mask intrinsic signals in the primary fibroblasts thus not allowing for inherent cell-cell cross talk to take place between stromal cells and tumor organoids. Of note, control organoids, which carried mutations in the *APC* gene, were grown in medium without supplementation of Wnt3a and R-spondin1 in order to enrich for tumor cells as previously described (*37*). Indeed, the viability and size of organoids co-cultured in the minimal medium with matched fibroblasts (O_CAF, O_NF) of three individual patients (P1-3) were comparable to these of organoids cultivated in regular organoid medium (O_ENAS) (Fig. 2A and fig. S2A, B), whereas organoid growth was not supported in minimal medium alone (O__min_). This indicated that factors derived from NFs or CAFs were equally potent to support organoid growth and survival as compared to conventionally used organoid medium.

**Fig. 2:**
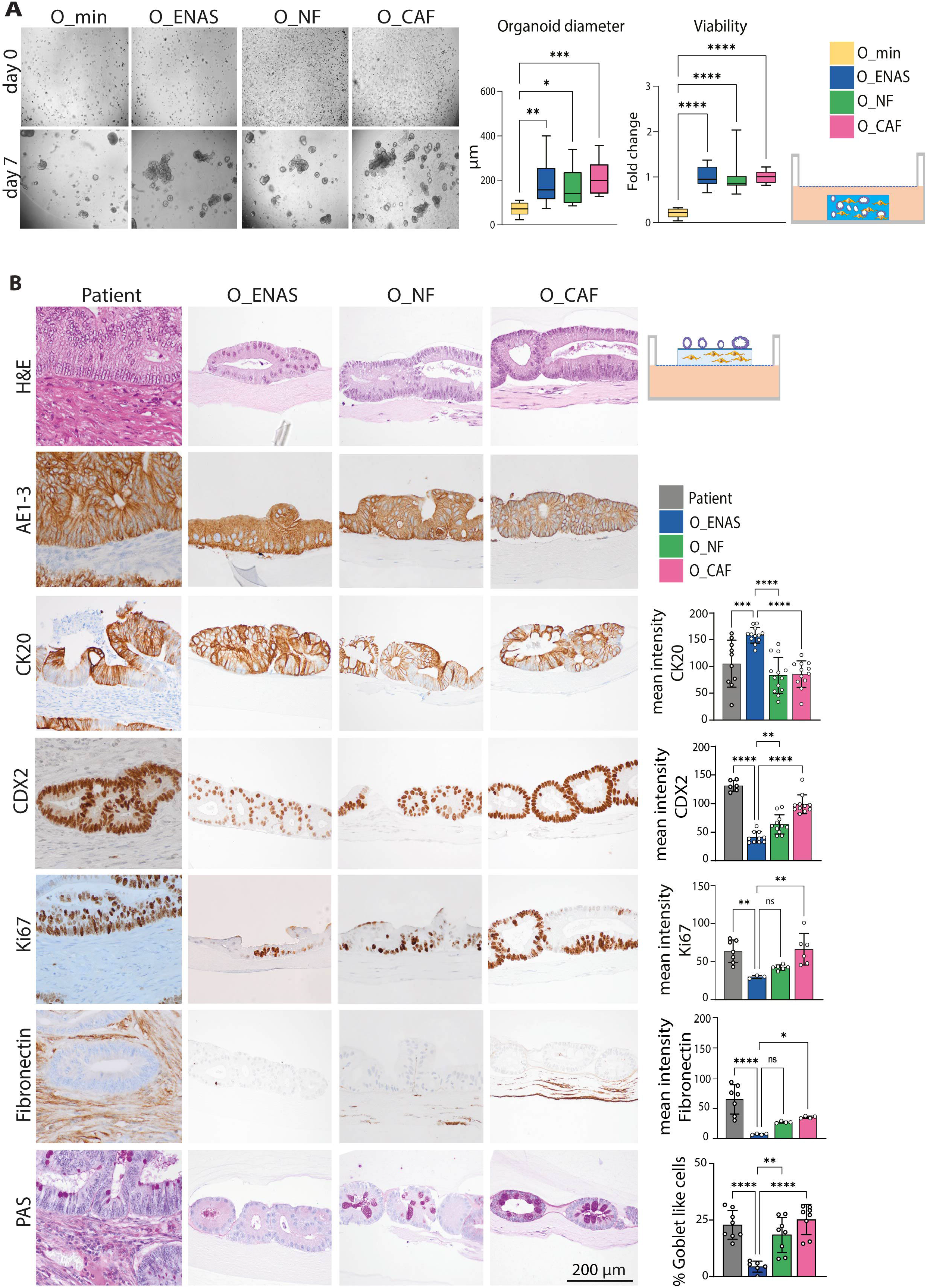
Organoid growth and morphology in co-cultures with fibroblasts. **(A)** Representative microscopic pictures of organoids grown in Matrigel for 7 days in minimal medium (O__min_), conventional organoid medium (O_ENAS) or together with NFs (O_NF) or CAFs (O_CAF) starting from single cell suspensions (left). On day 7 organoid diameters were measured using Fiji imaging software and viability was analyzed using Cell Titer Glo^®^ 3D assay (graphs on the right) (*N* = 3). The schematic on the right illustrates the co-culture setup in Matrigel. **(B)** Comparison of the histo-morphology between matched patient material and organoids grown either alone in ENAS medium or in co-culture with NFs or CAFs in minimal medium. Illustration on the top right displays the experimental setup. After cultivation for 7 days, the matrices were formalin fixed and paraffin embedded and sectioned into 2 μm thick slices. Consecutive slices were stained with haematoxylin and eosin (H&E), Periodic acid-Schiff stain (PAS) or via immunohistochemistry with indicated antibodies (AE1-3, pankeratin; CK20, cytokeratin 20; CDX2, differentiation marker; Ki67, proliferation marker; Fibronectin, fibroblast marker). Graphs on the right represent quantification of the images using Fiji software. A minimum of three images were quantified for each graph. Values presented are means ± SD, \**P* < 0.05, \*\**P* < 0.005, \*\*\**P* < 0.001, \*\*\*\**P* < 0.0001. Ordinary one-way ANOVA.

### Organoid-fibroblast co-cultures closely represent patient histology

Next, we evaluated whether a fibroblast/organoid co-culture model recapitulated the histological properties of the primary patient tumor. For this purpose, we embedded fibroblasts in a collagen I gel, which was coated with Matrigel and plated organoids on top. We used matched organoid/fibroblasts from three individual patients (P1-P3). This setup allows for direct cell-cell interaction and mirrors closely the *in vivo* tissue architecture of the colon. We used specific staining methods and immunohistochemistry such as hematoxylin and eosin (H&E) and periodic acid–Schiff (PAS) staining as well as immunohistochemical stainings with antibodies against CK20, CDX2, Ki67 and pan CK (AE1-3) (Fig. 2B and fig. S2C, D). We compared the histomorphology of the original tumor specimen to the one of NF and CAF co-cultures and organoid mono-cultures, grown in conventional organoid media. The co-cultures of the organoids with NFs and CAFs showed comparable morphology, which was highly similar to the patient material. Organoids grown in ENAS medium appeared slightly different to the co-cultured ones, displaying a more homogenous expression of certain markers, including CK20 and Ki67. Particularly, the heterogeneity in expression of the intestinal cell differentiation marker CK20 in the patient samples, was very closely recapitulated in the fibroblast co-cultures (fig. S2E). Thus, the epithelial organoid structures in the CAF and NF co-cultures, displayed the same distribution of undifferentiated, intermediate and highly differentiated cells as in the patient sample. In contrast, the organoids in ENAS medium displayed less similarity to the primary tumor and showed a more even CK20 expression based on quantification of positively stained cells (fig S2E). As expected, fibronectin, a marker of connective tissue cells and CAFs, was clearly present in the fibroblast layer of the co-cultures and in the stroma of the primary tumor but not in the ENAS organoid cultures in all of the three patients evaluated. Strikingly, PAS staining revealed that the number of mucus producing cells in the patient tumors was similar to the one in the *in vitro* co-cultures. However, mucus production was less pronounced in the organoids grown in full organoid medium without fibroblasts. In one of the tumors mucus was evident without the presence of clearly distinguishable mucus producing goblet like cells, which was also reflected in the co-cultures (fig. S2C). These histological data suggest that organoids cocultured with fibroblasts, both NFs or CAFs, manifest a significant similarity to the original *in vivo* tumor as indicated by the presence of various cell types.

### RNA sequencing reveals majorly deregulated genes between organoids grown with fibroblasts and ENAS medium

One exceptional advantage of the *in vitro* co-culture system, besides its physiological relevance, is the modularity of the system. The different components can be cultured individually or in combination, allowing to identify molecules, pathways and markers, which are selectively induced by the interaction of the tumor epithelium with the connective tissue cells. Thus, in order to dissect the crosstalk between the two components, we co-cultured organoids with patient matched fibroblasts for four days in a 3D setting and compared it to the single cultures of organoids or fibroblasts. In a slightly modified setup, organoids were cultured in Matrigel domes and fibroblasts in collagen I gels, however, the gels were positioned on the opposite sides of a transwell membrane insert allowing separate cultivation in the presence of soluble factor exchange and direct cell to cell interaction via the 0.45 μm diameter membrane pores as previously shown (*38*). After separation, the individual gels were lysed and mRNA prepared to perform bulk RNA sequencing (RNA-seq). Again, we used the same three patients as above and compared organoids in ENAS condition to organoids in combination with NFs or CAFs and organoids cultivated in minimal medium (O__min_), which was also used in the co-cultures. The high purity of the different cell populations (e.g epithelial cells and fibroblasts) was confirmed by selective expression of cell type specific genes including *CDH1, EPCAM* and *KRT8* for tumor cells and *VIM, DCN, THY1* and *COL3A1* for fibroblasts (fig. S3A, table S3). Interestingly, principal component analysis (PCA) of the transcriptomes of organoids grown in the different conditions showed a clear separation of organoids grown in ENAS or minimal medium from the organoid/fibroblast co-cultures (fig. S3B). Among the organoid fraction from the different conditions, we identified a total of 1742 significantly deregulated genes from pairwise comparison of the four different conditions (table S4). Supervised clustering of these deregulated genes revealed separate gene expression groups (Fig. 3A). The first group included genes upregulated in the O__min_ and ENAS conditions but downregulated in the cocultures, which were involved in MAPK1/MAPK3 signaling, PI3K/AKT signaling as well as signaling by interleukins. The second group of genes was specifically upregulated in the ENAS condition and contained genes involved in EGFR and FGFR signaling as well as cell cycle regulation genes. In the third group of genes, we observed genes upregulated in both co-culture conditions and downregulated in the ENAS and O__min_ conditions. This large group contained genes involved in extracellular matrix organization, collagen synthesis and modifying enzymes, as well as genes involved in innate immune system pathways. Finally, genes upregulated solely in the O__min_ condition, involved metabolism and differentiation associated genes. Reactome pathway analysis based on all 1742 significantly deregulated genes identified top altered pathways related to collagen biosynthesis and extracellular matrix organization (Fig. 3B). Interestingly several of the identified pathways were related to MET and FGFR signaling and cell motility. Thus, co-cultivation of fibroblasts with tumor organoids is strongly affecting ECM remodeling and cell-cell communication pathways in the tumor cells.

**Fig. 3:**
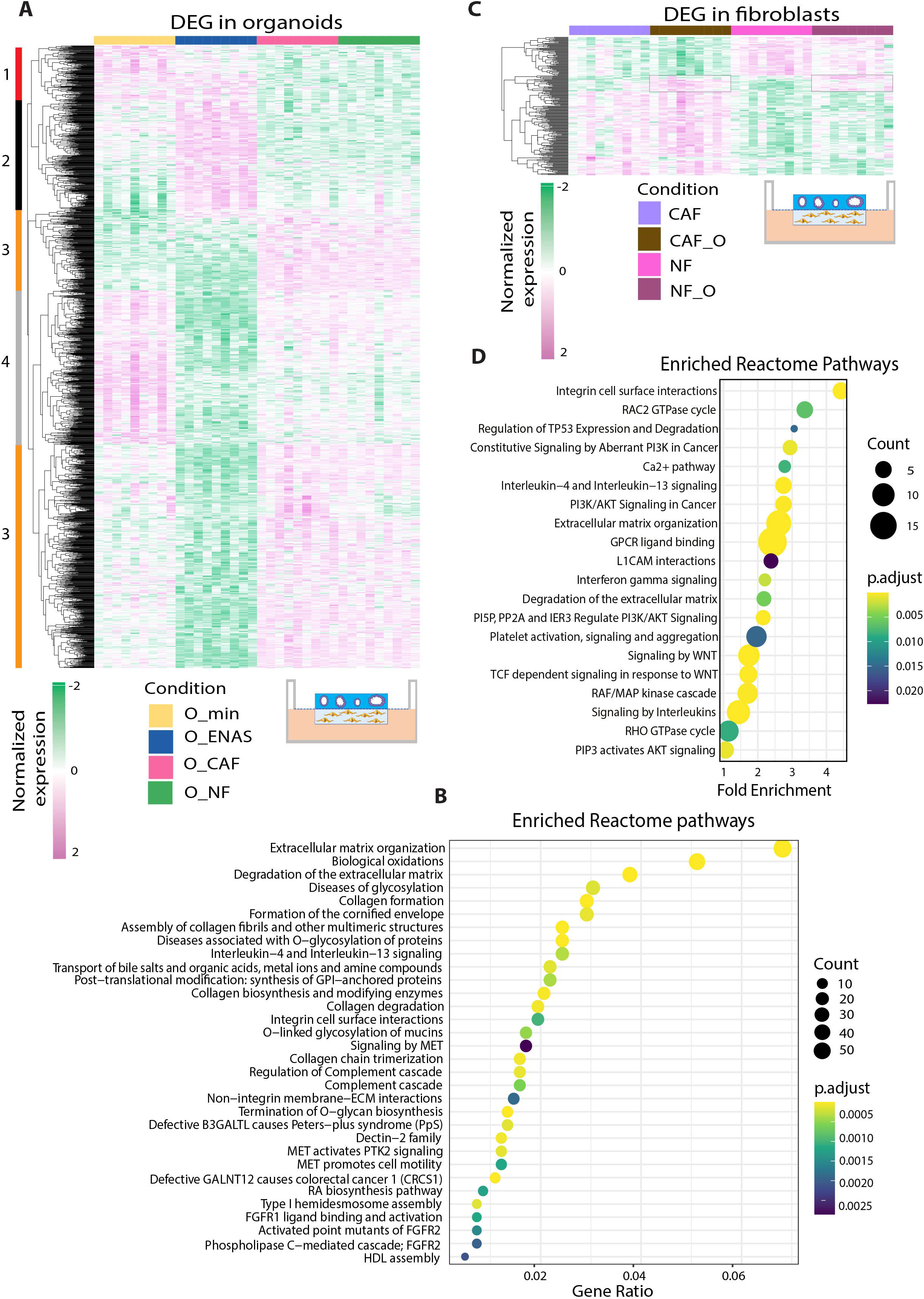
Gene expression analysis of tumor organoids and fibroblasts. **(A)** Hierarchical clustering of significantly differentially expressed genes in organoids grown in minimal medium (O__min_), conventional organoid medium (O_ENAS) or together with NFs (O_NF) or CAFs (O_CAF) as identified from RNA-seq analyses. Numbers and colors on the left of the heatmap indicate 4 different groups of genes that were upregulated in O__min_ (1, red), upregulated in O_ENAS (2, black), upregulated in the co-cultures O_NF and O_CAF (3, orange) or downregulated in O_ENAS (4, grey). The illustration below the heatmap shows the experimental setup. **(B)** Reactome pathway analysis shows the top significantly altered pathways between the individual growth conditions of organoids. Size of circles defines the number of genes affected in the respective pathway and the color indicates the significance (adjusted *P* value) according to the color gradient shown. **(C)** Hierarchical clustering of significantly differentially expressed genes of NFs and CAFs grown either alone (NF, CAF) or in co-culture with tumor organoids (NF_O, CAF_O). Rectangles on the heatmap mark genes that were specifically upregulated in the co-culture conditions in NFs and CAFs compared to the mono-cultures. **(D)** Bubble chart lists the top significantly changed pathways using Reactome pathway analysis in fibroblasts based on the different culturing conditions. The number of genes involved in the respective pathway is shown by the size of the circles and the significance by the color according to the color gradient. The RNA-seq analyses were performed in triplicates for three individual patients (P1-3) (9 replicates in total) (*P* adj. < 0.05).

### Co-culturing of NFs and CAFs with tumor organoids induces expression of specific genes

Next, we focused on the fibroblast expression profiles that were either cultured separately or in co-culture with the organoids. Using PCA analysis we found an overall separation between NFs and CAFs, with no clear segregation of NFs and CAFs cultured with organoids (fig. S3C). Similarly, hierarchical clustering revealed that the majority of differentially expressed genes allowed for the separation of NF versus CAF gene expression independent of culture condition (mono- or co-culture) (Fig. 3C). We identified 229 significantly differentially expressed genes by pairwise comparison of the four conditions (log2FC ≥1 *P* adj.<0.05) (table S5). Within the top differentially expressed genes between NFs and CAFs we found *WNT2, WNT5a, VCAM1* and *WISP1* upregulated in CAFs. These genes were previously identified as bona fide CAF markers *in vivo* and *in vitro* (*33, 39–41*). In NFs we found *RSPO3, TMEM35A* and *C7* enriched. *RSPO3* can potentiate the Wnt/β-catenin pathway independent of LGR binding (*42*). *TMEM35* codes for a transmembrane protein 35 that is preserved across species (*43*), however its exact function and particularly its role in cancer is poorly understood. The gene *C7* encodes for complement factor 7 and belongs to a family of proteins involved in immune surveillance and homeostasis. Additionally, C7 was identified as a tumor suppressor in ovarian cancer and non-small cell lung cancer (*44*). Reactome pathway analysis was comparable to results obtained from our proteomics analyses and revealed that a majority of molecules deregulated between NFs and CAFs belong to extracellular matrix rearrangement, cell motility and WNT signaling (Fig. 3D). Interestingly, also cancer pathways such as PI3K/AKT signaling and immune relevant pathways such as interleukin signaling were significantly enriched. Interestingly, we identified a group of 20 genes, which were specifically induced in both NFs and CAFs upon co-culture with organoids in all three patients analyzed (Fig. 3C and table S5). These genes including *IL6, ICAM-1* and *CXCL14*, have been associated with immune cell signaling, tumor progression and metastasis, indicating that tumor cells induce a highly specific gene expression signature to enforce their tumorigenic properties in the TME.

### Organoids co-cultured with fibroblasts manifest comparable gene expression to primary tumors

To validate the organoid-fibroblast model for its relevance in primary human tumors, we compared the expression levels of signature genes identified in our set of data to two recently published single cell RNA sequencing data sets published by Li *et al* (*14*) and Lee *et al*. (*13*) where human tumors were sequenced and cells categorized into various cell types and differentiation states based on their gene expression profiles. We inferred cell type specific marker genes from the two published datasets, which included markers for epithelial cells and fibroblasts and analyzed their expression in the bulk RNA-seq datasets from mono- and co-cultured organoids and fibroblasts (Fig. 4). The specific epithelial markers we evaluated were identified in stem cells, mature colonocytes, secretory cells (goblet cells) in the single cell analyses. We could determine four different gene groups: 1) genes overexpressed in 2) genes overexpressed in ENAS, and 3) genes upregulated in the co-cultures compared to the mono-cultures (Fig. 4A and table S6). As expected from the previous analyses, organoids grown in minimal medium (O__min_) showed a strong upregulation of differentiation specific genes such as *CLDN7, CDH1, KRT20* or *VIL1*. Also, genes specific for secretion such as *REG4*, *TFF1* and *TFF2* were upregulated in this condition. Genes specifically upregulated in the ENAS condition included *STMN1, CDCA7, LGR5* all of which are stem cell specific, in agreement with the stem cell supporting properties of the organoid expansion medium. In organoids co-cultured with fibroblasts, we identified stem cell, goblet cell, tuft cell and enterocyte specific markers including *RGMB, CEACAM1, TFF3, MUC1, SPINK1*, *CD44*, *OLM4*, *SPINK4* and *PTPRO*. Interestingly, there was no significant difference between genes expressed in organoids grown with NFs or CAFs except for *GUCA2B* and *FABP1*, which were slightly upregulated in the CAF condition. Overall, these data confirm the histo-morphological analyses indicating a shift from a stem-cell like to a more heterogenous pattern of cells in organoid co-cultures, which might be closer related to the *in vivo* tumor situation.

**Fig. 4:**
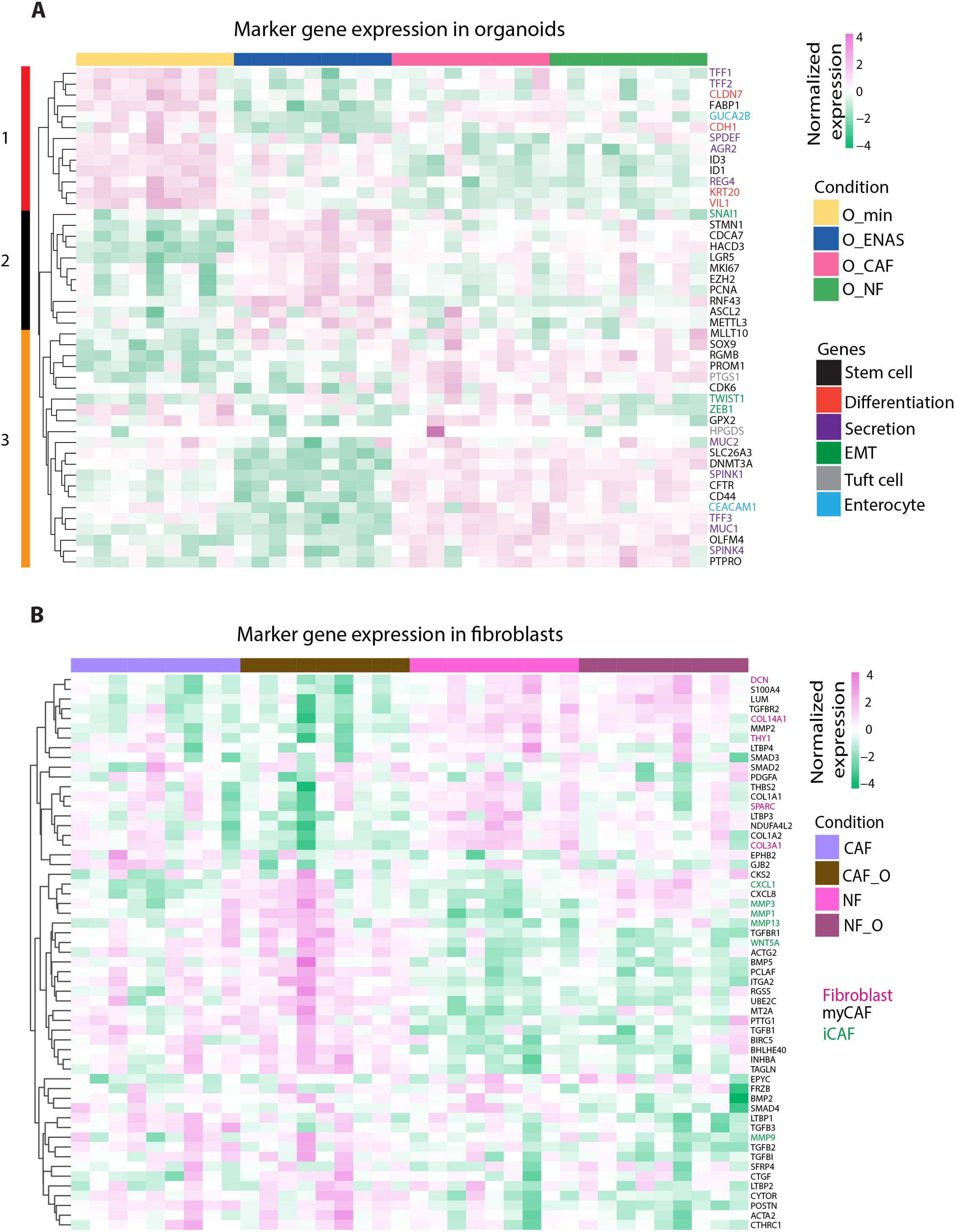
Expression of epithelial and fibroblast marker genes. **(A)** Hierarchical clustering of normalized gene expression levels of individual marker genes (derived from published scRNA-seq datasets of primary tumors (*13,14*) in organoids grown in minimal medium (O__min_), conventional organoid medium (O_ENAS) or together with NFs (O_NF) or CAFs (O_CAF). Colored rectangles and numbers on the left indicate genes upregulated in O__min_ (1, red), O_ENAS (2, black) or in the co-cultures with NF and CAF (3, orange). **(B)** Hierarchical clustering of general fibroblast, myofibroblast (myCAF) and inflammatory fibroblast (iCAF) markers (described in published scRNA-seq datasets of patient tumors (*13, 14, 25*) in cultured NFs and CAFs grown alone (NF, CAF) or in co-culture with organoids (O_NF, O_CAF).

### Fibroblast marker gene expression in co-culture with organoids reflects specific fibroblast subtypes

For the fibroblast markers, we derived marker genes identified in different subpopulations of the tumors including fibroblast and myofibroblast cells (*13, 14, 25*). We analyzed the expression of these genes in the bulk RNA-seq data from NFs and CAFs grown in the different conditions (Fig. 4B and table S7). As described above, major differences in fibroblast gene expression could be observed between NF and CAF independent of the culture condition (mono vs co-culture). However, we also identified several genes including *WNT5a*, *PCLAF*, *TAGLN, TGFB1* and *CXCL1* that were significantly upregulated in CAFs upon co-culture with the organoids. In fact, WNT5a has been previously reported as highly expressed in CAFs (*45*) and has a role in cancer progression. Additionally, *CXCL1, CXCL8, MMP1, MMP3* and *MMP13* were present at higher levels in the co-culture conditions (both NF and CAF), representing genes inducing proinflammatory and ECM degrading phenotypes in fibroblasts. These genes were defined as iCAF markers in scRNA-seq analysis (*25*).

Interestingly, some of the genes involved in collagen organization including decorin (*DCN*), which has been associated with tumor suppressive activities, and some collagen genes were downregulated upon co-culture with organoids in both NFs and CAFs. Also, Thrombospondin 2 (*THBS2*), a mediator of cell-cell and cell-matrix interactions, which suppresses angiogenesis and tumor growth was downregulated in the co-culture conditions. Together these data indicate that tumor organoids impact on the fibroblast phenotype by either enforcing CAF function or inducing pro-tumorigenic gene expression of NFs, and at the same time suppressing tumor inhibitory signaling.

### Deconvolution of bulk RNA sequencing data reveals distinct gene signatures in organoids grown in different conditions

To further validate the co-culture model, we employed a bioinformatics deconvolution strategy of our RNA-seq data by an unbiased approach using scRNA-seq analyses of primary CRC specimen from two published datasets from Lee *et al* (*13*) and Qian *et al* (*46*). This strategy allowed for the calculation of the proportions of specific cell types present in the individual culture conditions.

Firstly, we inferred marker genes from the Qian dataset to quantify cellular proportions of epithelial and fibroblast cells in the different culture conditions. The analyses showed a clear enrichment of fibroblast cells in the fibroblast subsets and a high specificity of epithelial cells in the organoid fractions of the RNA data (fig. S4A), confirming the purity of the cultures as already demonstrated with a small subset of specific marker genes above (fig. S3A). Gene expression levels of all markers identified for the deconvolution display a clear separation of fibroblasts and epithelial organoid cells using hierarchical clustering, as expected (fig. S4B and table S8).

In order to define the cellular proportions of organoids grown in the four different conditions, we combined tumor specific markers defining the consensus molecular subtypes (CMS1-4) and colon epithelial markers reflecting cellular subtypes and differentiation states from the Lee dataset (*13*) (table S9). The selected markers allowed for the subclustering of different epithelial cell states including stem-like transit amplifying (TA) and intermediate cells, goblet cells and mature enterocytes from the original dataset (*13*) (fig. S5). As previously described by Lee and colleagues, tumor epithelial cells are mainly represented by CMS2 and CMS3 gene expression profiles. Along these lines, we also observed enrichment of these subtypes in the organoid fractions (Fig 5A). Although we could observe interpatient heterogeneity among the organoid lines derived from the three individual patients, the ENAS condition mainly reflected the CMS2/3 tumor signatures. Upon cultivation in minimal medium (O__min_) a small proportion of differentiated mature enterocytes (type I and II) were apparent. Interestingly, co-culture with fibroblasts resulted in an increase of different cell type proportions, which was variable among the three patients but included also stem-like/TA and intermediate cells, goblet cells and mature enterocytes. Hierarchical clustering of marker gene expression levels of organoids in the different conditions revealed patient specific gene expression signatures, however, we also detected changes in marker gene expression based on the different conditions (Fig. 5B and table S9). In two of the three patients we identified a cluster of genes upregulated in both co-culture conditions including *MUC5AC*, *IFI6* and 27, *SerpinA1* and *LYPD8*, which were previously reported to be involved in tumor progression and metastasis, tumor cell proliferation and poor patient prognosis (*47–49*). In the ENAS condition we found upregulation of *NMU, EREG* and *SNHG12* genes. *NMU* and *EREG* encode for Neuromedin U and Epiregulin, respectively, both of which are involved in tumor invasion and progression (*50, 51*). *SNHG12* encodes for a long non-coding RNA, which can regulate tumor cell invasion and metastases in endometrial cancer (*52*). On the other side, a cluster of genes related to differentiation were downregulated in the ENAS condition, these genes include *GUCA2A*, *GUCA2B* and *KRT20*.

**Fig. 5:**
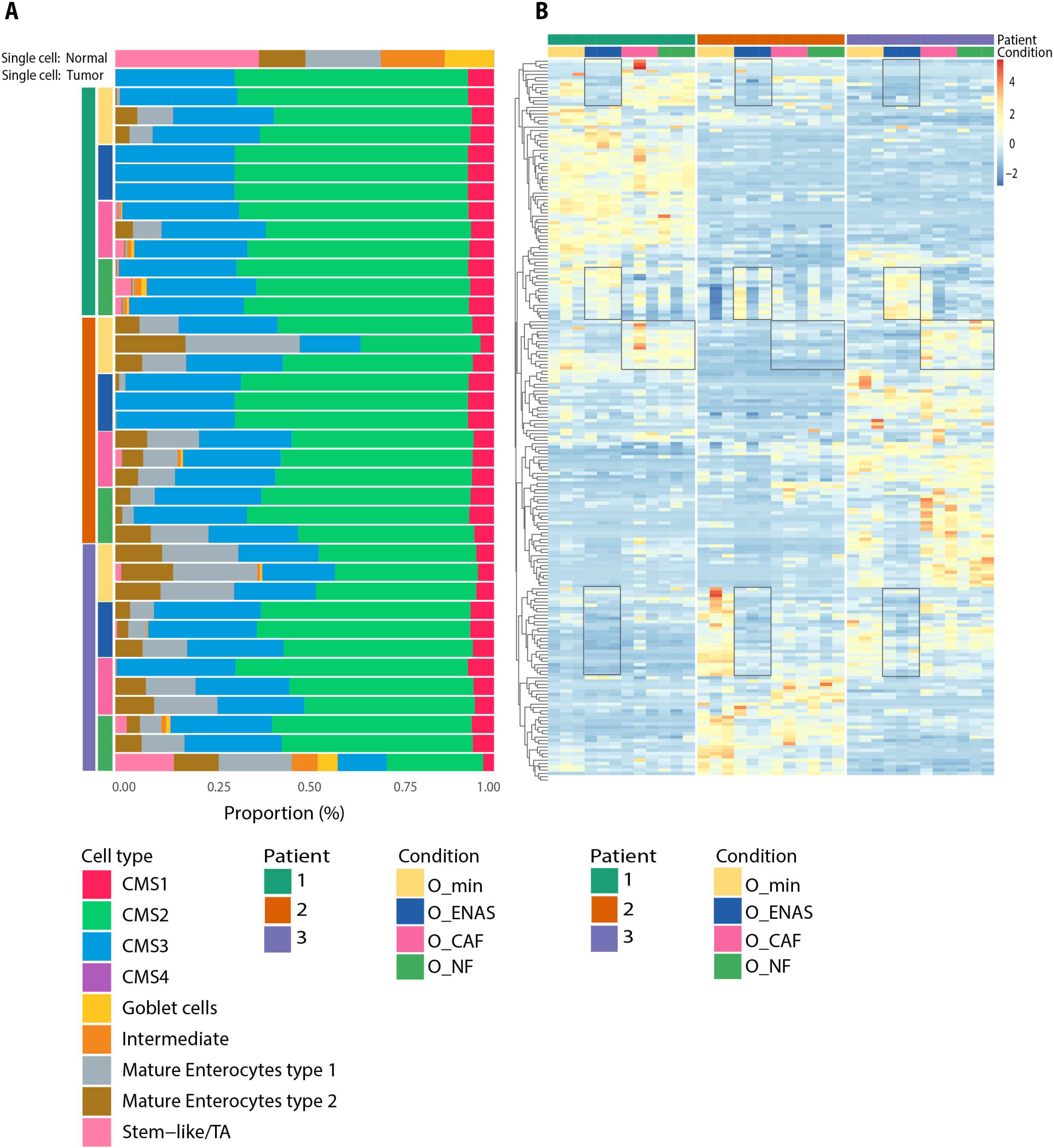
Identification of epithelial cell proportions in organoids. **(A)** Cellular proportions of organoids grown in minimal medium (O__min_), conventional organoid medium (O_ENAS) or together with NFs (O_NF) or CAFs (O_CAF), identified from deconvoluting bulk RNA-Seq data using scRNA-seq of 23 patient samples (13 tumor, 10 normal) from (*13*) as a reference. We derived marker genes for normal epithelial cells including 5 sub-types (stem-like/TA, goblet cells, intermediate, mature enterocytes type 1, and mature enterocytes type 2) and from tumor cells reflecting the four CMS signatures (CMS1-4). The top two rows in the graph display the cellular proportions of epithelial cells identified in primary normal tissues and tumors from (*13*), respectively. Subsequent rows represent the individual epithelial and CMS proportions of the organoids grown in the respective conditions, sorted by patient (P1-3). Different colors represent CMS1-4 and epithelial cell sub-types. Three replicates are shown for each condition. Please note that the last row, showing a replicate of organoids grown in co-culture with NFs, has a very high proportion of epithelial-like cells, which might represent an outlier. **(B)** Heatmap of gene normalized expression values of marker genes used for the deconvolution of organoid cell proportions as described in (A). For each patient (P1-3) and condition 3 replicates are shown. Rectangles mark gene expression signatures of marker genes that are deregulated in organoids grown in ENAS compared to other conditions, or in co-cultures (O_NF, O_CAF) compared to mono-cultures.

Thus, these data suggest that the co-cultivation of colon PDOs and fibroblasts induces a larger diversity of epithelial cell types as compared to the conventional system, where organoids are grown in monoculture and depend on stem cell niche factors.

## DISCUSSION

Organoid cancer models have shown great potential for precision medicine and there are living biobanks available, which are rapidly growing (*37, 53, 54*). They show predictive power for patient responses to some of the standard of care therapies such as irinotecan-based therapies in CRC (*5*). However, these models fail to predict patient response to oxaliplatin based therapies in CRC for unknown reasons. One reason might be the lack of the tumor stroma in these models, which evidently is also strongly affected by e.g. chemotherapy (*55*) and radiotherapy (*25, 56–58*). Here we present and thoroughly characterize a CRC PDO model integrating fibroblasts as a highly abundant and important cell type in the TME. We provide substantial evidence that this advanced model is mimics the biological and physiological properties of *in vivo* tumor conditions.

First, the preservation of the CAF phenotype under *in vitro* culture is still a matter of debate, and thus is the use of these cultured cells in cancer models. By comparing early passage cultured NFs and CAFs from the same patients we clearly demonstrate that bona fide CAF markers, which were identified *in vivo*, are preserved in culture. These include well characterized tumor stroma genes such as *WNT2* (*39*), *WISP1* (*59, 60*), *ITGB2* (*61*), *WNT5A* (*45*), *VCAM1* (*40*), *ICAM1* (*62*). Moreover, by transcriptomic and proteomic approaches as well as secretome analysis, we provide a list of differentially expressed molecules that are implicated in ECM organization as well as tumor and immune cell signaling. These molecular alterations were also apparent in the substantially elevated capacity of invasion/motility of CAFs into collagen gels. In line with our findings, CAFs have been described to display increased expression of matrix modifying enzymes and show increased remodeling capacity (*63, 64*), which is well recapitulated in the expression data of the cultured NFs and CAFs patient pairs, presented here.

However, fibroblasts serve not only as remodelers of the matrix, but they also have multiple signaling functions in the crosstalk with the surrounding cells supporting the development, spatial organization and homeostasis of epithelial sheets in many organs (*65*). Indeed, we could recapitulate the supportive nature of stromal fibroblasts for organoid growth *in vitro* and demonstrate that both NFs and CAFs grown in collagen I gels are equally potent in facilitating the survival and expansion of organoid structures of epithelial tumor cells. Of note, this cell culture system relies on minimal organoid medium supplemented only with 1% FBS omitting all protein factors (e.g. WNT3A, RSPO, Noggin, B27) and inhibitors (A-83-01, SB212090) otherwise required for monoculture organoid expansion in basal membrane extract or Matrigel (*28, 66*). This is of special importance for an undisturbed establishment of the physiological crosstalk between the fibroblasts and the epithelial cells. For example, it is well established that TGFβR inhibition (by e.g. A-83-01, and Noggin) critically affects fibroblast biology (*67, 68*) thus potentially masking physiologically relevant CAF phenotypes and crosstalk with the tumor cells. Such co-culture systems, albeit with synchronous cocultures from the start have been recently described in mouse and human and show similar results (*69–71*).

Noteworthy, the independent establishment of the stromal and organoid cultures in this system, provides one important intrinsic advantage, the possibility to analyze individual mono-cultures as well as the respective co-cultures including the mutual crosstalk of tumor cells and fibroblasts. Among the induced genes in both NFs and CAFs upon co-culture we found *IL6*, *ICAM1*, and *CXCL14*. IL6 is a cytokine which has been shown to induce epithelial to mesenchymal transition in gastric cancer cells (*72*). Intercellular adhesion molecule 1 (ICAM-1) has been implicated in several cancers including breast (*73*) and metastatic melanoma (*74*) by promoting cell invasion and transmigration, respectively. ICAM-1 was found to bind to MUC1 at the tumor leading edge thus promoting cell invasion (*75*). CXCL14 was found elevated in colorectal (*76*), ovarian (*77*), prostate (*78*) and breast (*79, 80*) cancers. Stromal overexpression of CXCL14 in ovarian cancer was linked to poor patient survival via STAT3 dependent manner due to increased tumor cell proliferation (*77*). Taken all together, these data suggest that tumor cells are instructing CAFs on the production of specific molecules, which then in turn allow tumor cells to propagate and invade. The experimental setup here was designed to co-culture the cells for 4 days, which possibly is only the starting point where tumor cells transform fibroblasts into a supportive tumor entity and longer cultures would possibly induce larger changes in the fibroblasts. On the other hand, the co-culturing conditions with either NFs or CAFs significantly upregulated genes in organoids related to extracellular matrix organization and collagen formation and biosynthesis, which suggests that phenotypic alterations are not limited only to fibroblasts.

Strong support for the physiological meaningfulness of the model presented here comes from histo-morpholgoical analyses of the co-cultures. The level of similarity of marker protein expression and presence of mucus was higher between the patient tissue and the co-cultures as compared to organoids alone – either in minimal or ENAS medium. Furthermore, mucus presence in tumors is usually related to worse patient outcome (*81*), and thus its presence might be relevant in cultures, when drug response is tested. Also, RNA seq results when juxtaposed to single cell sequencing data from CRC supported the presence of various cell types in the fibroblast co-cultures as observed in *ex vivo* tumors (*13, 14*). This demonstrates that organoid expansion medium is indeed enriching stem cells as expected (*28*), whereas the heterogeneity of cell types is inferior to that in organoids grown with fibroblasts exemplified by e.g. the presence of goblet cell markers or tuft cell markers in this group. Organoids in minimal medium showed an increase in differentiation markers such as *KRT20, CLDN7, VIL* (*82–84*) as anticipated but failed to show diversity. Thus, the culture system presented here shows high levels of tumor cell heterogeneity, which is important for cancer progression and drug response and could hence be of important translational impact (*85–87*).

Deconvolution of the bulk RNA sequencing data of this study demonstrated expression of genes found in primary tumors. Particularly the genes *MUC5AC*, *IFI6* and *IFI27*, *SerpinA1*, *LYPD8*, *CHP2* were predominantly expressed in organoids co-cultured with fibroblasts but not in monocultures. As these genes are majorly found to promote tumor growth and metastases formation, these data agree with previous reports that fibroblasts enhance tumor progression (*88–90*). Furthermore, this demonstrates that there is considerable difference in expression patterns in PDOs when they are grown as a mono-versus co-culture with fibroblasts. These differences need to be considered when performing drug screens with specific inhibitors or searching for altered signaling pathways in an *in vitro* system.

Taken together, we established a system, which closely recapitulates the primary tumor biology in its morphology and gene expression patterns and allows studying relevant tumor-stroma communication. Future work would be needed to dissect expression changes within single cells in both the tumor and the stromal compartment, ideally involving the full repertoire of cells including the fibroblasts, immune cells and possibly endothelial cells. Additionally, it will be important to validate this culture system for its accuracy to recapitulate patients’ responses to standard of care therapy. This will provide more precise treatment options and allow for the stratification of patients to optimized therapy solutions. In addition, we foresee high potential of this system to screen for new therapeutic options, which target both tumor cells and the surrounding stroma, which might help to limit therapy resistance and result in better treatment outcome.

## MATERIALS AND METHODS

### Study design

The aim of this study was to generate a PDO model from CRC tumor specimen including stromal fibroblast cells isolated from the same patient. For this purpose, we used primary patient tumors or adjacent normal mucosa to establish and characterize organoids and fibroblasts – both cancer associated or normal. The in-depth molecular characterization of the model was done in comparison to classical organoid mono-cultures and a control condition, using three individual patient samples, representing three biological replicates. All experiments were further done in at least three independent replicates.

### Ethics approval for the patient material

Patient material was obtained upon signed consent and the experiments were performed according to the “Good Scientific Practice Guidelines” of the Medical University of Vienna, as well as the latest “Declaration of Helsinki”. The ethical protocol has been reviewed and approved by the ethics review board of the Medical University of Vienna (Votum N# 1248/2015).

### Organoid and fibroblast isolation

Tumor pieces were processed either fresh or viably frozen. Every piece was washed three times with PBS and then minced into 1 mm pieces. Pieces were transferred into a 15 ml tube and incubated in 5 ml DMEM 10 % FBS and 1 mg/ml collagenase type I at 37 degrees for 1 hour. Then the digest was applied onto a 70 μm strainer. The single cell suspension was washed three times with PBS as in the last wash the pellet was divided into two – one half was embedded into Matrigel on a 24 well plate and the other half was plated on a 10 cm dish with EGM2-MV (PromoCell Cat# C22022) medium for fibroblasts growth.

### Organoid and fibroblast culture

Organoids were cultured in 30 μl Matrigel droplets in 24 well plates and passaged every 3 to 4 days. They were grown in organoid medium as previously described (*28*). In order to enrich for tumor cells Wnt3a and Rspondin 1 were omitted from the medium. After isolation, fibroblasts were grown in EGM2-MV (PromoCell Cat# C22022) medium and passaged every 3 to 4 days in T75 flasks.

### Mutational profile of organoids

The mutational profile of the organoids and patients was evaluated using the Ion AmpliSeq Cancer Hot Spot Panel v2 (Thermo Fisher, Cat# 4475346), which includes the following genes: *ABL1, EGFR, GNAS, KRAS, PTPN11, AKT1, ERBB2, GNAQ, MET, RB1, ALK, ERBB4, HNF1A, MLH1, RET, APC, EZH2, HRAS, MPL, SMAD4, ATM, FBXW7, IDH1, NOTCH1, SMARCB1, BRAF, FGFR1, JAK2, NPM1, SMO, CDH1, FGFR2, JAK3, NRAS, SRC, CDKN2A, FGFR3, IDH2, PDGFRA, STK11, CSF1R. FLT3, KDR, PIK3CA, TP53, CTNNB1, GNA11, KIT, PTEN, VHL*.

### IHC staining and quantification

For immunohistochemical stainings co-cultures were prepared. Fibroblasts were embedded in collagen gels as described. When gels had solidified, they were coated with 10 μl Matrigel for 20 minutes and lifted on cell culture inserts, where the Matrigel coating was upwards. On top we plated organoids at 50 organoids per gel. To each well 1.5 ml medium was added underneath to ensure the organoids were grown at the air-liquid interphase. The co-cultures were grown for 1 week and then fixed in 4% paraformaldehyde for 20 minutes. Then they were embedded in paraffin and sectioned at 2 μm sections for staining. Antibodies used were anti-CDX-2 (Biogenex Cat#NC1701134), anti-Ki67 (Ventana Medical Cat#790-4286), anti-CK20 (Ventana Medical Cat# 790-4431) and anti-Fibronectin (Dako Diagnostics Cat#A0245). Staining intensity was quantified using Fiji software as described earlier (*91*).

### Viability assay

For assessing the viability of organoids and fibroblasts, the cells were plated in 10 μl Matrigel drops on a 96 well plate. Each drop contained either fibroblasts alone (10^4^ cells per well) or organoids (2×10^3^ cells per well) or both. To each well 90 μl medium was added and medium was exchanged every other day. After 7 days of culturing 100 μl of CellTiter-Glo 3D (Promega Cat# G9681) was added to each well and incubated for 5 minutes on a shaker. Luminescence was measured using Synergy™ HTX Multi-Mode Microplate Reader.

### Sprouting assay

NFs and CAFs were trypsinized and counted. For formation of spheroids, 500 cells were plated per well in U bottom 96 well plates in 100 μl DMEM 5% FBS/Methylcellulose 5%. Plates were incubated for 5 h in the incubator to allow spheroids to form. Then spheroids were collected, washed with PBS and embedded into collagen gels from rattail collagen I at 2mg/ml. When the gels were solidified, they were moved to 24 well plates and spheroids were grown for 24 h and imaged thereafter. Quantification was performed using Fiji software by drawing the outgrowth area of each spheroid and subtracting the middle spheroid area; single cells and number of protrusions were counted separately. The data was plotted in GraphPad Prism.

### Cytokine arrays

For evaluating the secretome of NFs and CAFs, 2×10^6^ cells were plated per T75 flask in EGM2-MV medium. The next day, the medium was exchanged with DMEM / 1% FBS and incubated for 48 h. Then, medium was collected and centrifuged for removal of floating cells. From that, 1 ml was used for further analysis. The assay was performed according to the manufacturer’s instructions (R&D systems Cat# ARY022B). Readout of fluorescence was performed on Odyssey DLx and the fluorescence intensity was quantified with Image Studio Lite Ver 5.2 software.

### RNA collection and sequencing

For RNA collection organoids and fibroblasts were grown as co-cultures onto inserts with a 0.45 μm membrane. Fibroblasts were embedded into collagen gels from rat tail collagen I at a concentration of 2 mg/ml. The gels were casted onto one side of the membrane and organoids were plated in 100 μl Matrigel drops on the other side of the membrane. The co-culture was grown in 6 well plates, where the fibroblast gels were positioned on the lower side and the organoids on the top. To each culture, 1.5 ml medium was added – either ENAS medium or 1% FBS in Advanced F12/DMEM and the cells were cultured for four days. After four days, the gels were lyzed and total RNA was collected using QIAGEN RNeasy kit (Cat# 74106). RNA quantity was evaluated with a Qubit™ Fluorometer and 200 ng RNA were used for bulk RNA sequencing.

Sequencing libraries were prepared using the NEBNext Ultra II Directional Kit with polyA selection module (New England Biolabs, MA, USA). In brief, total RNA was used as an input into the mRNA selection with oligo-dT paramagnetic beads. mRNA was then fragmented and converted into double stranded cDNA. Following universal adapter ligation, samples were PCR barcoded using dual indexing primers. Samples were sequenced in depth of 30 to 52 million paired-end 2×150bp reads on Illumina NovaSeq 6000 sequencer (Illumina, CA, USA).

### Bioinformatic analysis of RNA-seq data

Bcl files were converted to Fastq format using bcl2fastq v. 2.20.0.422 Illumina software for base calling. Quality check of raw paired-end fastq reads was carried out by FastQC (http://www.bioinformatics.babraham.ac.uk/projects/fastqc/). The adapters and quality trimming of raw fastq reads was performed using Trimmomatic v0.36 (*92*) with settings CROP:250 LEADING:3 TRAILING:3 SLIDINGWINDOW:4:5 MINLEN:35. Trimmed RNA-Seq reads were mapped against the human genome (hg38) and Ensembl GRCh38 v.94 annotation using STAR v2.7.3a (*93*) as splice-aware short read aligner and default parameters except --outFilterMismatchNoverLmax 0.1 and --twopassMode Basic. Quality control after alignment concerning the number and percentage of uniquely- and multi-mapped reads, rRNA contamination, mapped regions, read coverage distribution, strand specificity, gene biotypes and PCR duplication was performed using several tools namely RSeQC v2.6.2 (*94*), Picard toolkit v2.18.27 (http://broadinstitute.github.io/picard) and Qualimap v.2.2.2 (*95*) and BioBloom tools v 2.3.4-6-g433f (*96*). The differential gene expression analysis was calculated based on the gene counts produced using RSEM tool v1.3.1 (*97*) and further analyzed by Bioconductor package DESeq2 v1.20.0 (*98*). Data generated by DESeq2 with independent filtering were selected for the differential gene expression analysis due to its conservative features and to avoid potential false positive results. Genes were considered as differentially expressed based on a cut-off of adjusted p-value ≤ 0.05 and log2(fold-change) ≥1 or ≤-1. Clustered heatmaps either for organoids or fibroblasts sample fractions were generated from selected top differentially regulated genes using the R package pheatmap v1.0.10 (*99*). The gene selection was performed separately for fibroblasts and organoids, where differential expression analysis was carried out for all possible condition pairs followed by union of significantly differentially expressed genes from all comparisons. All heatmaps were based on logarithmic DESeq2 normalized genes expressions with removed patient batch effect and further normalized on Z-score. We utilized hierarchical clustering to sort genes into individual clusters. Volcano plots were produced using ggplot v3.3.3 package (https://doi.org/10.1002/wics.147) and MA plots were generated using ggpubr v0.4.0 package version 0.1 7 (2018) (https://CRAN.R-project.org/package=dply.). PCA plots were based on DESeq2 normalized gene expression, further adjusted by variance stabilizing transformation as well as by removing patient batch effect and utilizing prcomp function from stats v3.6.3 package (https://www.R-project.org/). Reactome pathway analyses including pathways significance bar and dot plots as well as network connection figures were executed by adapting of clusterProfiler v3.12.0 package (*100*).

### Proteomics

Proteins were precipitated with four volumes ice-cold acetone at −20°C overnight. After centrifugation at 10000 x g for 30 min the supernatant was discarded and the remaining pellets washed with 150 μL 80% ice-cold acetone. After centrifugation at 10000 x g for 5 min the supernatant was discarded and remaining acetone allowed to evaporate. The protein pellet was resuspended in 50 μL 8 M urea in 50 mM ammonium bicarbonate and the protein concentration determined using a Bradford assay (BioRad). Proteins were reduced using 10 mM dithiothreitol for 30 min at room temperature, alkylated with 20 mM iodoacetamide for 30 min at room temperature in the dark, and remaining iodoacetamide quenched with addition of 5 mM dithiothreitol. Aliquots of 60 μg protein were diluted to 4 M urea using 50 mM ammoniumbicarbonate, then 600 ng Lys-C were added and the samples incubated at 25°C for 2 h. After dilution to 1 M urea 600 ng trypsin were added and the samples incubated at 37°C overnight. The samples were acidified with 0.5 % trifluoroacetic acid and one fifth of each sample was desalted using C18 Stagetips (*101*). Sample quality and concentration was determined by injecting 3% of each sample on a HPLC-UV system at 214 nm, using a monolithic C18 column for peptide separation.

An estimated 600 ng peptides were separated on an Ultimate 3000 RSLC nano-flow chromatography system (Thermo-Fisher), using a pre-column for sample loading (Acclaim PepMap C18, 2 cm × 0.1 mm, 5 μm, Thermo-Fisher), and a C18 analytical column (Acclaim PepMap C18, 50 cm × 0.75 mm, 2 μm, Thermo-Fisher), applying a segmented linear gradient from 2% to 35% and finally 80% solvent B (80 % acetonitrile, 0.1 % formic acid; solvent A 0.1 % formic acid) at a flow rate of 230 nL/min over 120 min. Eluting peptides were analyzed on a Q Exactive HF-X Orbitrap mass spectrometer (Thermo Fisher), which was coupled to the column with a nano-spray ion-source using coated emitter tips (PepSep, MSWil).

The mass spectrometer was operated in data-dependent acquisition mode (DDA), survey scans were obtained in a mass range of 375-1500 m/z with lock mass activated, at a resolution of 120k at 200 m/z and an AGC target value of 3E6. The 25 most intense ions were selected with an isolation width of 1.2 m/z, fragmented in the HCD cell at 28% collision energy and the spectra recorded for max. 54 ms at a target value of 1E5 and a resolution of 15k. Peptides with a charge of +2 to +6 were included for fragmentation, the peptide match and the exclude isotopes features enabled, and selected precursors were dynamically excluded from repeated sampling for 30 seconds.

Raw data were processed using the MaxQuant software package version 1.6.0.16 as described (*102*) and the Uniprot human reference proteome (www.uniprot.org, release 2018_08), as well as a database of most common contaminants. The search was performed with full trypsin specificity and a maximum of two missed cleavages at a protein and peptide spectrum match false discovery rate of 1%. Carbamidomethylation of cysteine residues were set as fixed, oxidation of methionine and N-terminal acetylation as variable modifications. Label-free quantification the “match between runs” feature and the LFQ function were activated - all other parameters were left at default.

MaxQuant outputs were filtered for potential contaminants, reverse hits and in case they were only identified by site. Additionally, only protein groups that were quantified in at least 75% of the samples in at least one condition (i.e. 3 out of 4 in CAF or NF) have been included for further downstream differential protein expression analysis using custom R-scripts mainly using functions from the DEP package (*103*). After intensity normalization by variance stabilizing transformation (*104*) and confirming left-censored nature of missing values, remaining missing values were imputed applying the MinProb method (*105*). Statistical testing for differentially expressed proteins was done by using functions implemented by the DEP package, making use of protein-wise linear models combined with empirical Bayes statistics (*103*). Secretome measurements were analysed for differential intensity similarly, missing values were not present.

Subsequent pathway enrichment analyses were performed via active subnetworks using the pathfindR package (*106*). As background protein-protein interaction network we selected STRING (*107*), as input only significantly differentially expressed proteins with a p-value lower than 0.05 and an absolute log2-fold-change higher than 1 (i.e. two-fold) and differentially secreted proteins with a p-value lower than 0.05 for secretome data were used.

### Statistics

Statistical analyses were performed with Prism software (GraphPad, Version 9). For the study either two-tailed Student’s *t* test or one-way analysis of variance (ANOVA) were used, as indicated. A *P* value <0.05 was set as a threshold of significance.

### Deconvolution

We use a modified version of the SCDC library (*108*) to calculate the proportions of the bulk RNA-seq with different scRNA-seq as a reference. The general procedure allows the deconvolution of bulk RNA-seq expression samples based on a scRNA-seq dataset with a multiple reference cell type clustering avoiding pre-defined marker genes. These are the steps in the procedure – first, a signature matrix is built based on the cell type scRNA-Seq reference. Next, the cell-type proportions for the bulk samples are recovered using a tree-guided procedure and a weighted non-negative least square (W-NNLS) regression framework over the marker genes, weighting each gene by cross-subject and cross cell variation. The method defines an observed bulk gene expression 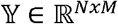 for N genes across M samples, each containing K cell types. The deconvolution constructs two non-negative matrices 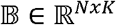(signature matrix) and 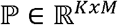(joint proportions of the K cell type by sample) such that:

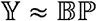

The SDCC library has been modified to allow parallelism, matrix sparsity, and selection of marker genes using a dynamic threshold for each cluster in the single-cell reference matrix. The native approach for the selection of marker genes was completed with a global threshold using a Wilcoxon test between each cell type and the remaining dataset. This creates an unbalanced set of marker genes, where cell types have either a very high number, few, or zero markers. To overcome this limitation, this has been substituted by a process with multiple bootstrapping samples on all clusters selecting n_b_ clusters as background and then performing a Wilcoxon test over each sampling and cluster of interest. Lastly, using DBSCAN to perform outlier analysis, the procedure selects an optimum number of genes that behave as markers for the specific cell type.

### Deconvolution of samples

We used 73 bulk data samples divided into two groups: 36 organoids and 37 fibroblasts. As a reference we used the following two scRNA-seq datasets: SMC dataset (*13*): This contains data from scRNA-seq of unsorted CRC single cells of 23 patients, out of which 13 are from tumor samples and 10 are from normal samples. The authors originally identified five cell types (stromal, Myeloid, Mast, T, B, and Epithelial cells) through clustering, and four consensus molecular subtypes (CMS1, CMS2, CMS3, and CMS4). After filtering just the epithelial cells, we recovered 18,539 cells. Of these, 1,070 corresponded to normal epithelial cells with five sub-cell types (Stem-like/TA, Goblet cells, Intermediate, Mature Enterocytes type 1, and Mature Enterocytes type 2) and 17,469 tumor cells with the four CMS clusters described earlier. In order to run the deconvolution with the tree-guided procedure, we divided the 9 cell types into the following groups: Cluster group 1: CMS1, CMS2, CMS3 and CMS4; Cluster group 2: Stem-like/TA, Goblet cells, Intermediate; Cluster group 3: Enterocytes type 1, Mature Enterocytes type 2. Using an initial pseudo bulk data simulated based on the single-cell subset, we determined the optimal parameters for deconvolution selecting n_b_ = 4 background cell clusters, selected n_bs_ = 100 bootstrapping samples, and limited the number of required marker genes per cluster to be between N_min_ = 28, N_max_= 35. With the optimal parameters, we obtain a Pearson correlation value of >0.999 between the true proportions and the estimated proportions and a sum of residuals < 3.6e-28. After the simulation, we ran the deconvolution with the actual bulk data and we obtained 220 marker genes and a deviance value of 8.97 (0.12 on average for each of the 73 samples). Pan-cancer blueprint single-cell profiling dataset (*46*): This dataset includes data about CRC colorectal cancer from 7 patients, 14 normal tissues, and 7 with cancer. The study includes 9 cell types clustered (epithelial, fibroblast, B cell, cancer, endothelial, enteric glia, mast cells, myeloid and T cell). After filtering, we retrieved four cell types: epithelial, fibroblast, cancer, endothelial defined by 44,684 cells, where 14,058 were normal epithelial cells with 5 cell types and 30,626 tumor cells. In order to run the deconvolution with the tree-guided procedure we grouped the 4 cell types as follows: Cluster group 1: epithelial, fibroblast; Cluster group 2: cancer, endothelial. Using a pseudo bulk data simulated based on the singlecell subset, we determined the optimal parameters for deconvolution selecting n_b_ = 2 background cell clusters, chosen n_bs_ = 100 bootstrapping samples and setting the number of required markers genes per cluster to be between N_min_ = 28, N_max_= 35. With the optimal parameters, we obtain a Pearson correlation value of >0.9059 between the estimated proportions and the true proportions and a sum of residuals < 340.90 (170,45 in average for normal and tumor samples). After the simulation, we executed the deconvolution with the actual bulk data and we obtained 137 markers genes and deviance of 7,774.67 (106.50 on average for each of the 73 samples. Finally, we used the marker genes to visualize the normalized expression values by sample.

## List of Supplementary Materials

Fig S1 to S5.

Data files S1 to 9 (Excel files)

## Acknowledgments

The authors thank the International Max Planck Research School for Intelligent Systems (IMPRS-IS) for supporting Crhistian de Jesus Cardona.

## Funding

Austrian Science Fund (FWF), doc.funds grant DOC59 (GE, TM)

Österreichische Forschungsförderungsgemeinschaft (FFG) 879481(GE, MB, VSA, JK)

Austrian Academy of Sciences, Doc Fellowship 25276 (LT)

CCC research grant of the MedUni Vienna (HD)

European Commission, SECRET ITN 859962 (HD)

Niederösterreichische Forschungs und Bildungsges.m.bH.; NFB, LSC18 017 (HD)

BMBF, German Network for Bioinformatics Infrastructure (de.NBI) 031A537B, 031A533A, 031A538A, 031A533B, 031A535A, 031A537C, 031A534A, 031A532B (CJC, GS)

## Author contributions

Conceptualization: GE, HD, VSA

Methodology: VSA, HD, GE, JK, CB, MHa, LT, TM

Pathological assessment and tissue acquisition: CB, KW, LM, MB

Investigation: VSA, BN, JC

Bioinformatics: CJC, VH, AT, MHa

Visualization: VSA, GE, HD, VH, CJC, AT

Funding acquisition: GE, MB, VSA, MM, HD, MHe

Project administration: VSA, GE, HD

Supervision: GE, HD, GS, BT

Writing – original draft: VSA, GE, HD, VH, CJC

Review: all authors

## Data and materials availability

### Source Code deconvolution

For more details, you can see a complete R notebook and the source code at: https://github.com/crhisto/CRC_organoids_and_fibroblasts_notebook

### Proteomics data

The mass spectrometry proteomics data have been deposited to the ProteomeXchange Consortium via the PRIDE partner repository (*109*) with the dataset identifier PXD.

### RNA-sec data

The RNA-seq data sets were deposited to the NCBI Gene Expression Omnibus (GEO) (*110*) accession number GSE.

