## Supplementary material for "Mimicking tumor cell heterogeneity of colorectal cancer in a patient-derived organoid-fibroblast model": Suplementary Figures

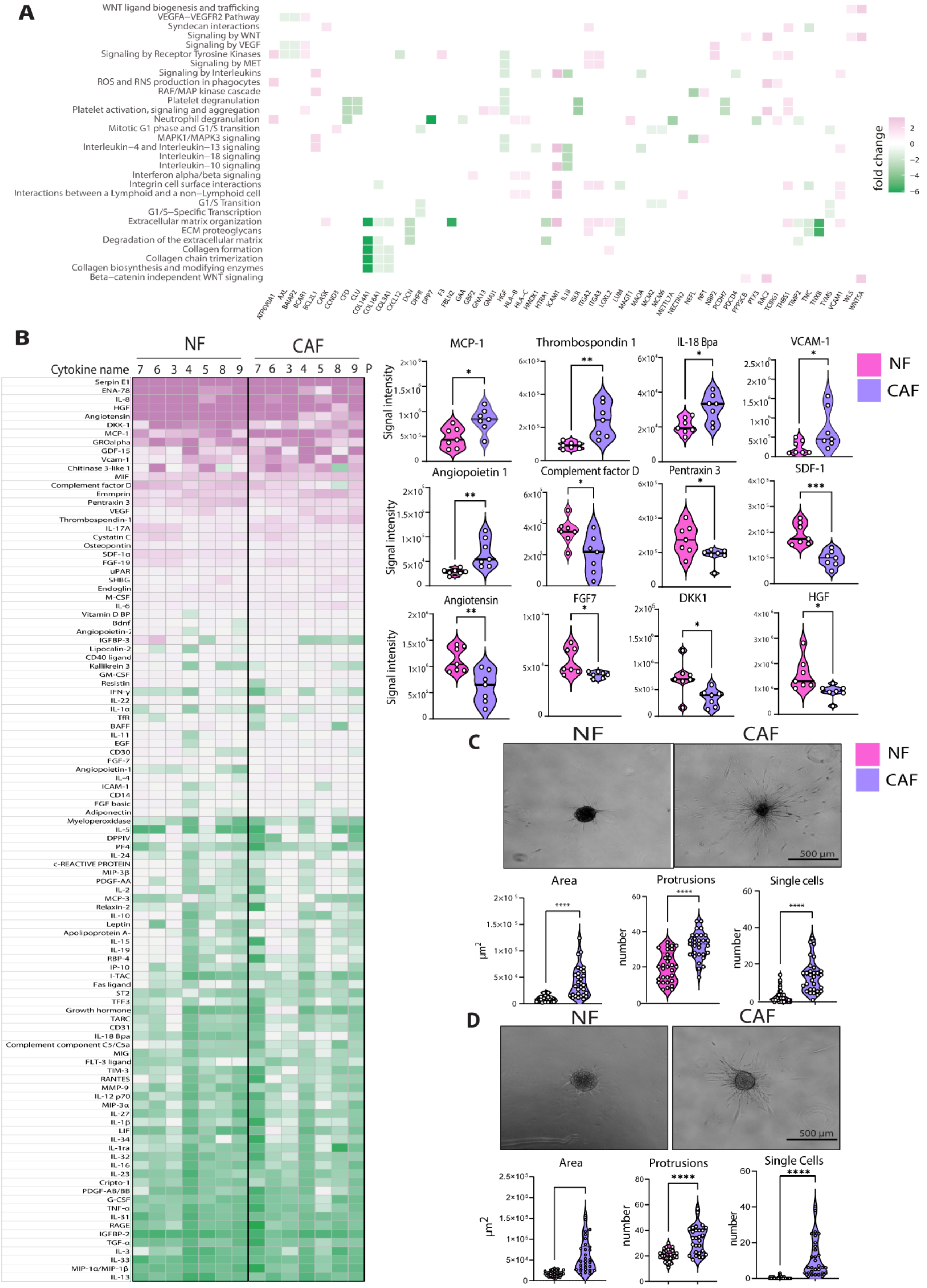
A
WNT ligand biogenesis and trafficking
VEGFA−VEGFR2 Pathway
Syndecan interactions
Signaling by WNT
Signaling by VEGF
Signaling by Receptor Tyrosine Kinases
Signaling by MET
Signaling by Interleukins
ROS and RNS production in phagocytes
RAF/MAP kinase cascade
Platelet degranulation
Platelet activation, signaling and aggregation
Neutrophil degranulation
Mitotic G1 phase and G1/S transition
MAPK1/MAPK3 signaling
Interleukin−4 and Interleukin−13 signaling
Interleukin−18 signaling
Interleukin−10 signaling
Interferon alpha/beta signaling
Integrin cell surface interactions
Interactions between a Lymphoid and a non−Lymphoid cell
G1/S Transition
G1/S−Specific Transcription
Extracellular matrix organization
ECM proteoglycans
Degradation of the extracellular matrix
Collagen formation
Collagen chain trimerization
Collagen biosynthesis and modifying enzymes
Beta−catenin independent WNT signaling
fold change
2
0
−2
−4
−6
B
NF
CAF
Cytokine name
7 6 3 4 5 8 9 7 6 3 4 5 8 9 P
MCP-1
Thrombospondin 1
IL-18 Bpa
VCAM-1
Angiopoietin 1
Complement factor D
Pentraxin 3
SDF-1
Angiotensin
FGF7
DKK1
HGF
Signal intensity
C
NF
CAF
500 µm
Area
Protrusions
Single cells
µm2
number
D
NF
CAF
500 µm
Area
Protrusions
Single Cells
µm2
number

**Fig. S1: Molecular and phenotypic differences of NFs and CAFs. (A)** Heatplot shows majorly deregulated pathways between matched NFs and CAFs of 4 patients and respective up- and downregulated proteins involved, derived from LC MS/MS analyses. **(B)** Intensity levels of 105 cytokines analyzed from conditioned media of cultured matched pairs of NFs and CAFs from 7 patients (left). Violin plots (right) show protein intensity levels of significantly differentially secreted proteins from the 7 individual NF and CAF pairs (*$P < 0.05$, **$P < 0.005$, unpaired two-tailed Student's *t* test) (right). **(C)** and **(D)** Representative microscopic pictures of NF and CAF spheroids embedded into collagen I for 24h (left). Violin plots show measurements of the sprouting area (total area minus spheroid core area), number of protrusions and number of single cells detached from the spheroids (right). Experiments were performed in triplicates and 10 individual spheroids were analyzed for each experiment ($N = 30$). *$P < 0.05$, **$P < 0.005$, ***$P < 0.001$, ****$P < 0.0001$, two tailed unpaired Student's *t* test).

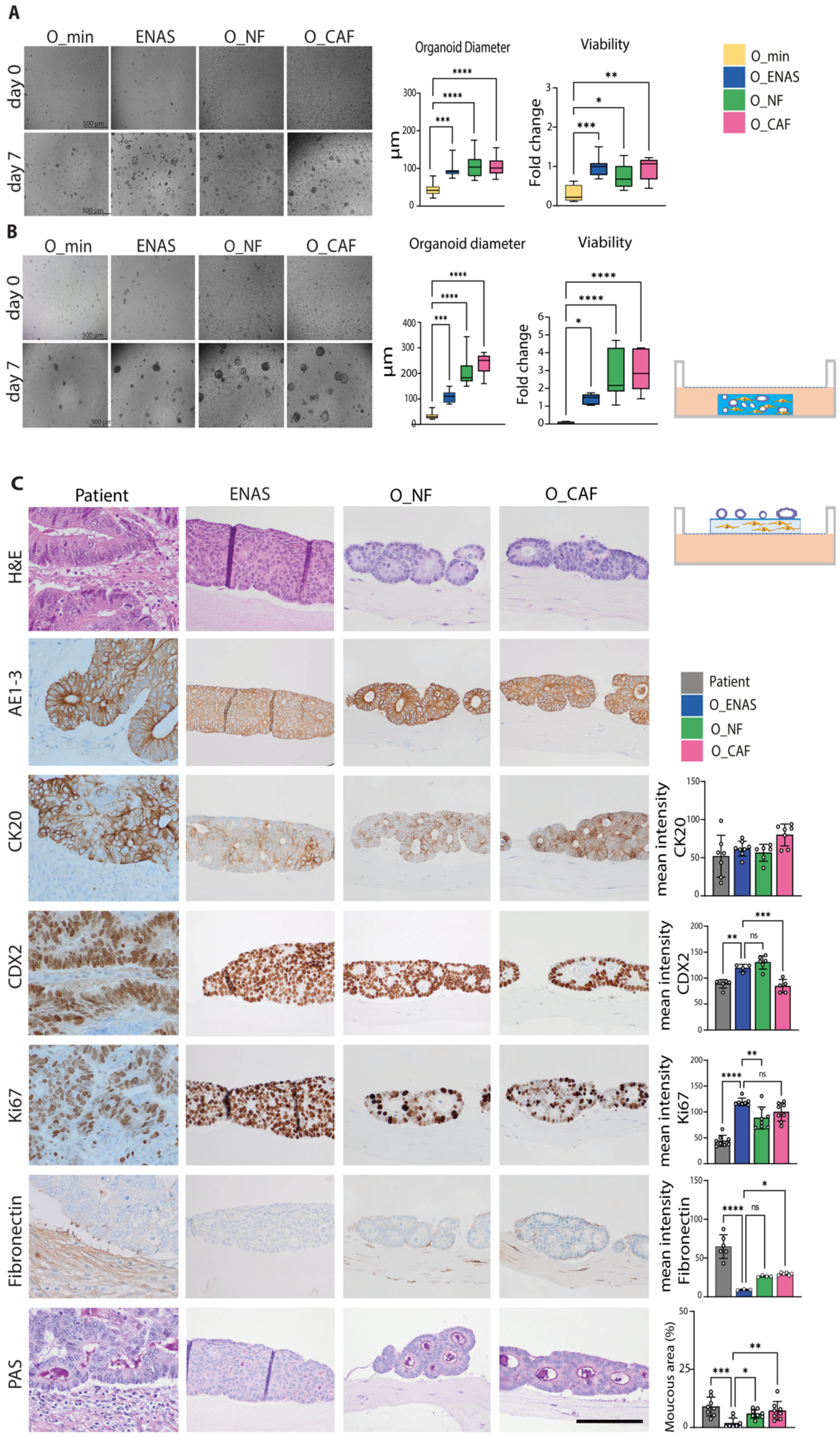
A
O_min
ENAS
O_NF
O_CAF
day 0
day 7
500 µm
Organoid Diameter
Viability
µm
Fold change
O_min
O_ENAS
O_NF
O_CAF
B
O_min
ENAS
O_NF
O_CAF
day 0
day 7
Organoid diameter
Viability
µm
Fold change
C
Patient
ENAS
O_NF
O_CAF
H&E
AE1-3
CK20
CDX2
Ki67
Fibronectin
PAS
Patient
O_ENAS
O_NF
O_CAF
mean intensity CK20
mean intensity CDX2
mean intensity Ki67
mean intencity Fibronectin
Moucous area (%)

D

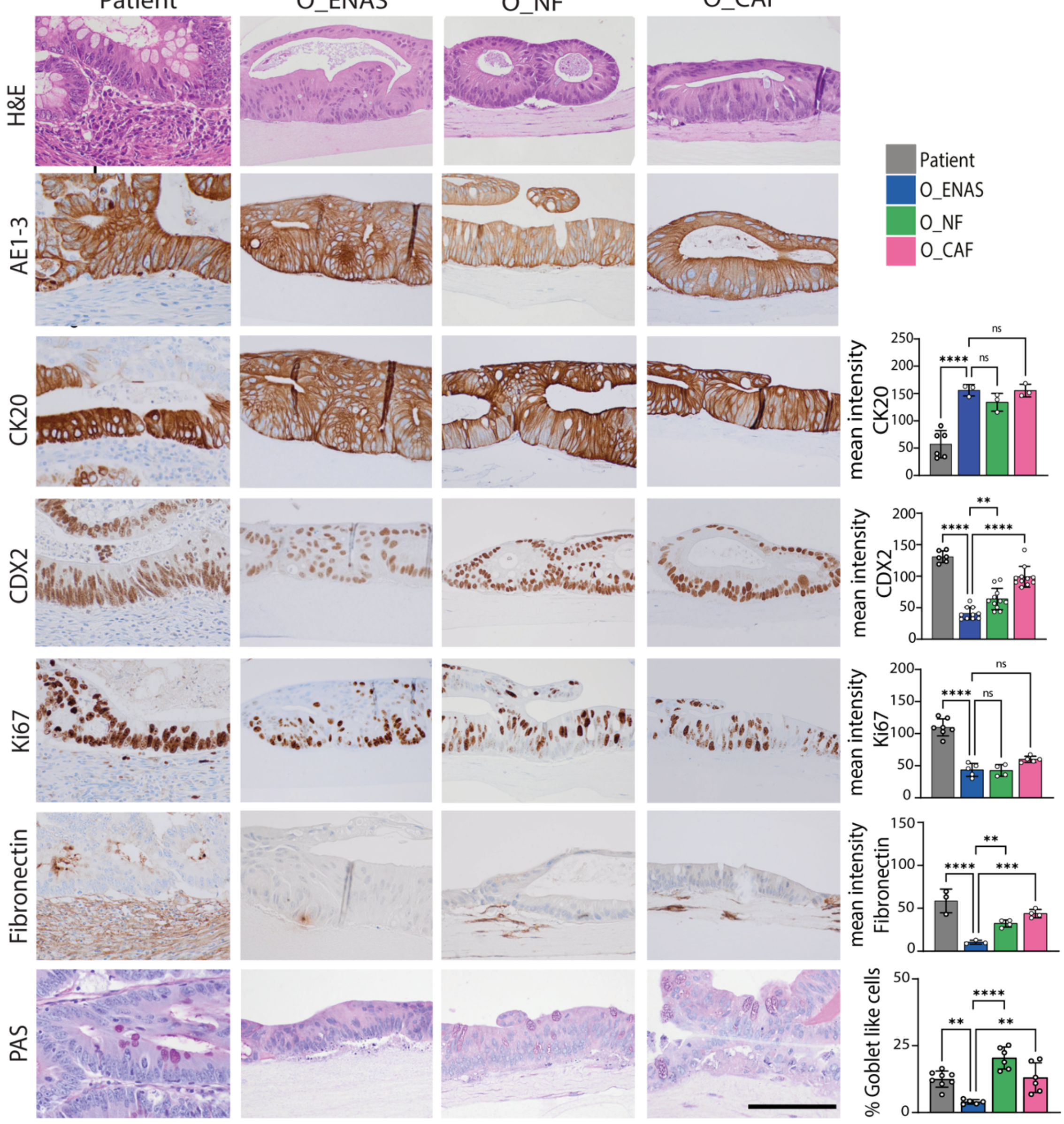
Patient
O_ENAS
O_NF
O_CAF
H&E
AE1-3
CK20
CDX2
Ki67
Fibronectin
PAS
Patient
O_ENAS
O_NF
O_CAF
mean intensity CK20
mean intensity CDX2
mean intensity Ki67
mean intencity Fibronectin
% Goblet like cells
ns
****
**
***

E

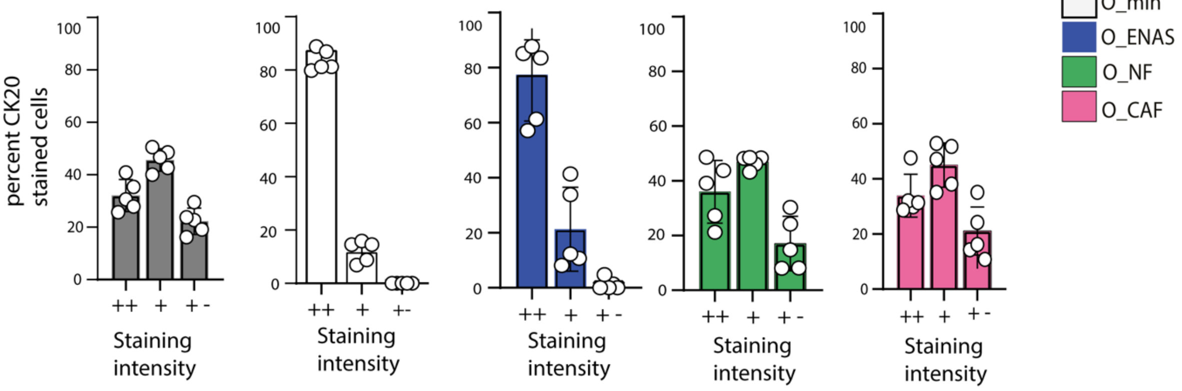
percent CK20 stained cells
Staining intensity
++
+
+-
Patient
O_min
O_ENAS
O_NF
O_CAF

**Fig. S2: Organoid growth and morphology in co-cultures with fibroblasts. (A)** and **(B)** Representative microscopic pictures of organoids grown in Matrigel for 7 days in minimal medium ($O\_{min}$), conventional organoid medium (O_ENAS) or together with NFs (O_NF) or CAFs (O_CAF) starting from single cell suspensions (left). On day 7 organoid diameters were measured using Fiji imaging software and viability was analyzed using Cell Titer Glo® 3D assay (graphs on the right) ($N = 3$). The schematic on the right illustrates the co-culture setup in Matrigel. **(C)** and **(D)** Comparison of the histo-morphology between matched patient material and organoids grown either alone in ENAS medium or in co-culture with NFs or CAFs in minimal medium. Illustration on the top right displays the experimental setup. After cultivation for 7 days, the matrices were formalin fixed and paraffin embedded and sectioned into 2 µm thick slices. Consecutive slices were stained with haematoxylin and eosin (H&E), Periodic acid-Schiff stain (PAS) or via immunohistochemistry with indicated antibodies (AE1-3, pan-keratin; CK20, cytokeratin 20; CDX2, differentiation marker; Ki67, proliferation marker; Fibronectin, fibroblast marker). Graphs on the right represent quantification of the images using Fiji software. A minimum of three images were quantified for each graph. Values presented are means ± SD, $*P < 0.05$, $**P < 0.005$, $***P < 0.001$, $****P < 0.0001$. Ordinary one-way ANOVA. **(E)** Quantification of intensity of CK20 staining for one patient shown in Fig. 2B, ++ high, + medium and + - low intensity. Data are mean values ±SD.

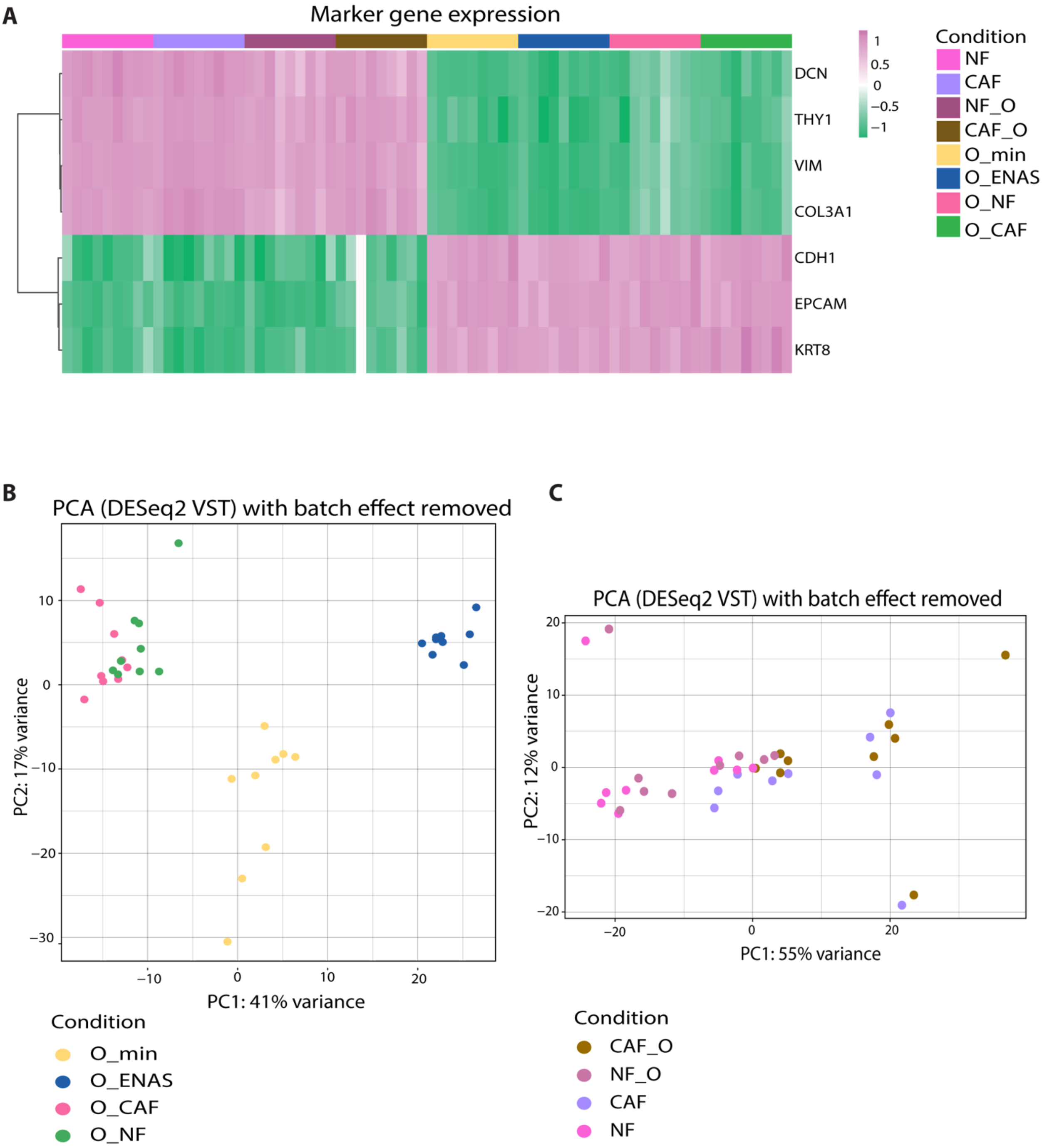


**Fig. S3: Gene expression analysis of organoids and fibroblasts. (A)** Hierarchical clustering of gene expression levels of classical cell type specific genes for tumor/epithelial and fibroblast cells isolated from mono- or co-cultures and analyzed by RNA-seq. **(B)** Principal component analysis (PCA) plot of organoid samples grown in the different conditions plotted in two dimensions using their projections onto the first two principal components based on RNA-seq data. The different conditions are color coded. **(C)** PCA plot of fibroblasts grown in the different conditions (indicated by colors) plotted in two dimensions using their projections onto the first two principal components based on RNA-seq data.

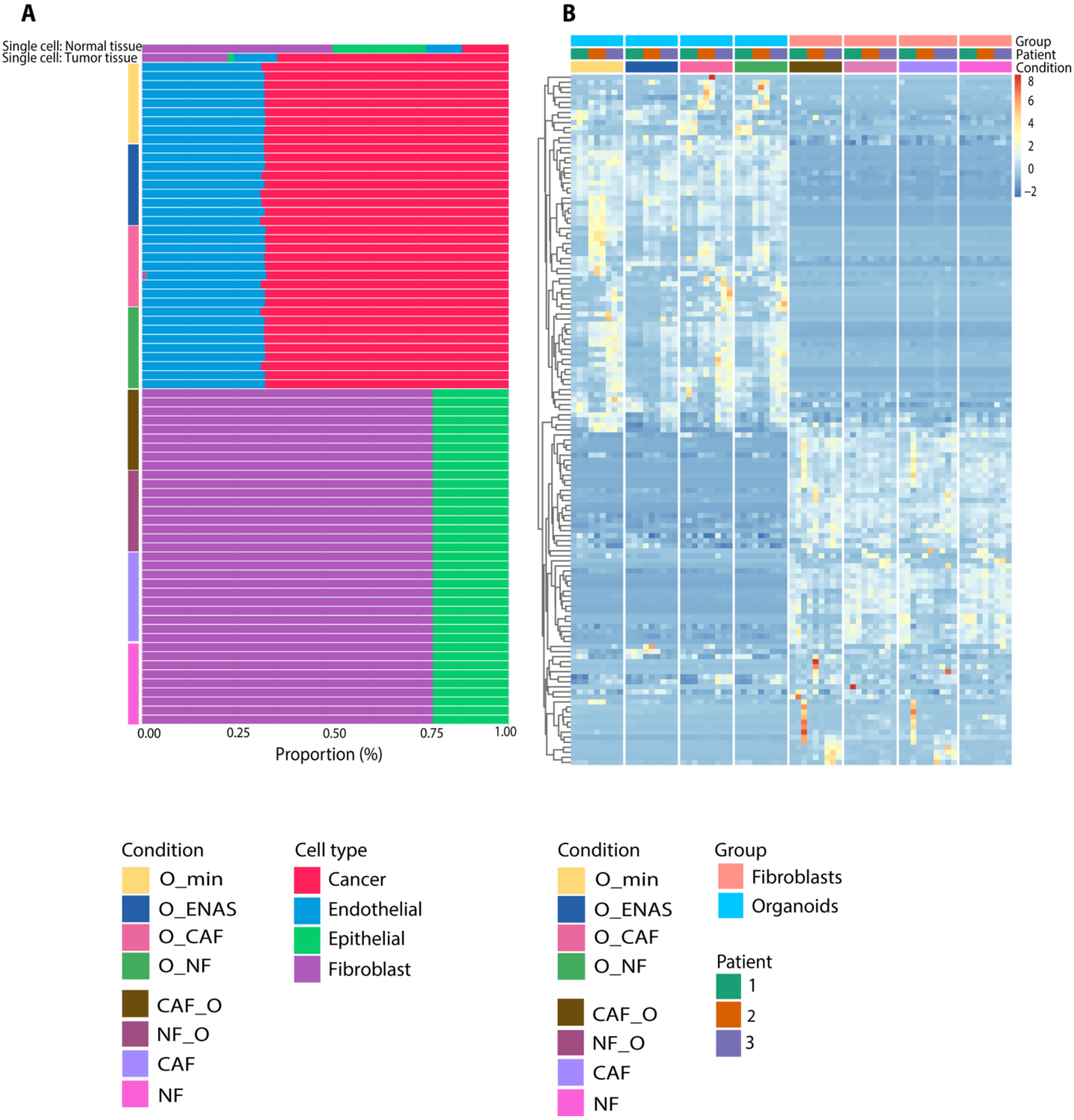


**Fig S4: Deconvolution of bulk RNA sequencing data based on scRNA-seq of primary CRC tumors. (A)** Validation of cell type purity of tumor and fibroblast cells from mono- or co-cultures by deconvolution based on 137 marker genes inferred from a published CRC scRNA-seq data set *(46)* **(B)** Hierarchical clustering of bulk RNA-seq gene expression levels of 137 marker genes used for the deconvolution in (A) in NFs, CAFs grown alone (NF, CAF) or in co-culture with organoids (NF_O, CAF_O).

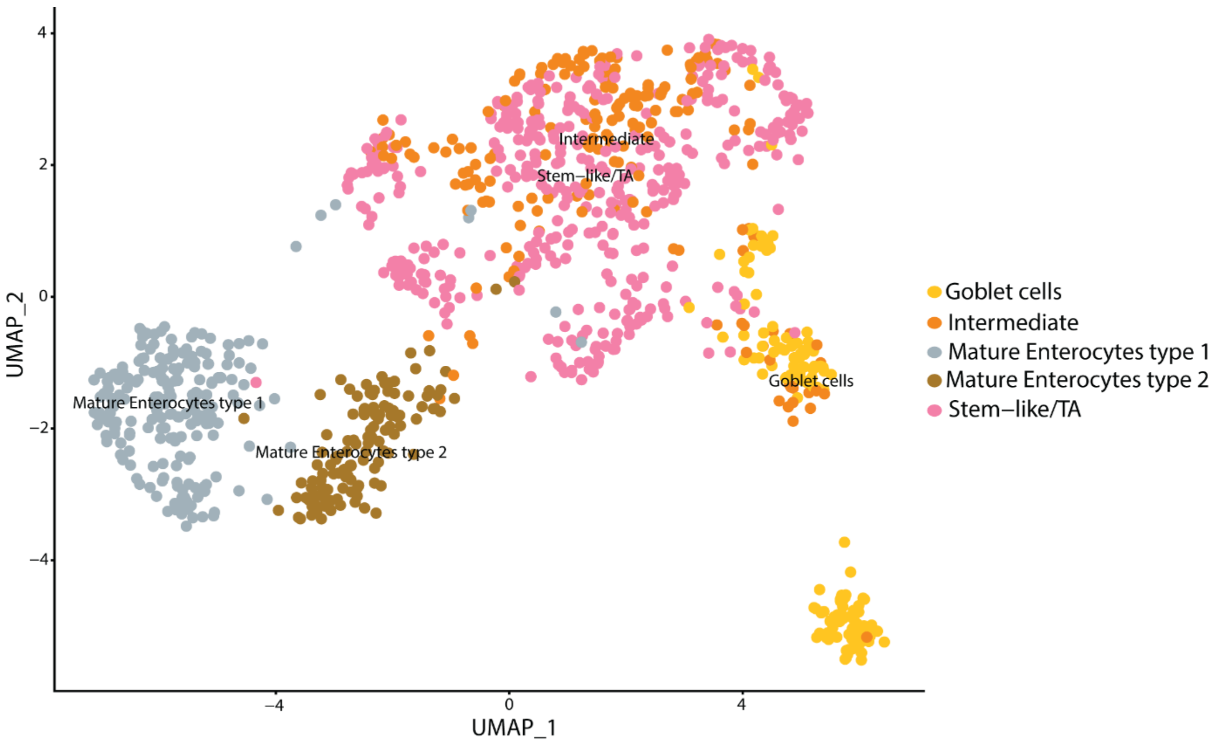


**Fig S5: Cell type identification.** UMAP representation of subclustering analysis based on epithelial marker genes identified in scRNA-seq datasets *(13)*. The gene set includes markers for goblet cells, intermediate stem cells, mature enterocytes type 1 and 2 and stem-like/transit amplifying (TA) cells.
